# Three Distinct Transcriptional Profiles of Monocytes Associate with Disease Activity in SSc Patients

**DOI:** 10.1101/2022.01.29.477568

**Authors:** Hadijat-Kubura M. Makinde, Julia L.M. Dunn, Gaurav Gadhvi, Mary Carns, Kathleen Aren, Anh H. Chung, Lutfiyya N. Muhammad, Jing Song, Carla M. Cuda, Salina Dominguez, John E. Pandolfino, Jane E. Dematte D’Amico, G. Scott Budinger, Shervin Assassi, Tracy Frech, Dinesh Khanna, Alex Shaeffer, Harris Perlman, Monique Hinchcliff, Deborah R. Winter

**Affiliations:** Northwestern University, Feinberg School of Medicine Department of Medicine, Division of Rheumatology. Chicago, IL 60611; Cincinnati Children’s Hospital Medical Center, Division of Allergy & Immunology. Cincinnati, OH 45229 (current affiliation).; Northwestern University, Feinberg School of Medicine Department of Preventive Medicine. Chicago, IL 60611; Northwestern University, Feinberg School of Medicine, Department of Medicine, Division of Gastroenterology and Hepatology. Chicago, IL 60611.; Northwestern University, Feinberg School of Medicine, Department of Medicine, Division of Division of Pulmonary and Critical Care. Chicago, IL 60611; Prospective Registry of Early Systemic Sclerosis (PRESS) consortium. Shervin Assassi MD MS- University of Texas at Houston (TX) Elana Bernstein MD MS- Columbia University (NY) Robyn Domsic MD MS - University of Pittsburgh (PA) Tracy Frech MD MS - University of Utah (UT) Jessica Gordon - Hospital for Special Surgery (NY) Faye Hant - Medical University of South Carolina (SC) Monique Hinchcliff – Yale School of Medicine (CT) Dinesh Khanna MD MS - University of Michigan (MI) Ami Shah - Johns Hopkins University (MD) Victoria Shanmugam - George Washington University (DC); University of Texas Health Science Center at Houston, Division of Rheumatology and Clinical Immunogenetics, Houston, Texas 77030.; Vanderbilt University, Department of Medicine, Division of Rheumatology and Immunology. Nashville, TN 37232; University of Michigan, Department of Medicine, Division of Rheumatology. Ann Arbor, MI 48109.; Yale School of Medicine Section of Rheumatology, Allergy & Immunology. New Haven, CT 06520

## Abstract

**Background/Purpose:** Patients with systemic sclerosis (SSc) display a complex clinical phenotype. There are numerous studies that relate transcriptional signatures from PBMC or whole skin of SSc patients to disease activity. However, analyses of whole tissue RNA-sequencing studies are subjected to changes in cellular composition that can drive gene expression signatures and a loss of the ability to detect biologically important transcriptional changes within minority cell populations. Here, we focused on circulating monocytes, which have been shown to exist as two central populations classical (CM) and non-classical (NCM).

**Methods:** SSc patients were recruited from four different sites that form PRESS: Northwestern University, University of Texas, University of Michigan and University of Utah. Comprehensive clinical data was collected for all patients. We isolated CM and NCM from these patients and age, sex, and race-matched healthy volunteers were used as controls. RNA-seq was performed on CM and NCM populations as well as on isolated bulk macrophages from skin.

**Results:** We first performed RNA-seq on CM, which are the predominant population in circulation. In order to capture the variability across the SSc cohort, we defined 1790 differentially expressed genes in each patient. We then used these genes to cluster patients into 3 subgroups: Groups A-C. Group A exhibited the strongest interferon signature and innate immune pathways. Group B patients expressed genes in the same pathways but was also enriched for response to cAMP and corticosteroids. Both Group B and Group C exhibited upregulation of genes associated with vasculature development and blood vessel formation. Group C uniquely upregulated TGFB pathways. Next, we performed RNA-seq on NCM isolated from the same patients. When NCM were clustered based on the same 1790 genes as CM, we found that Groups A and C were recapitulated, while Group B was less cohesive. Our analysis stratified SSc patients based on their transcriptional profiles in monocytes but was agnostic to their clinical presentation. We found that Group B and C patients exhibited significantly worsened lung function at the time of monocyte isolation than Group A patients. However, there were no significant differences in skin disease. We then isolated macrophages from skin biopsies of SSc patients and showed that the transcriptional profile of Group A and C in SSc patients was conserved. We also used gene expression data from another study on monocytes which stratified patients based on disease presentation. We found that Group A accurately distinguished dcSSc and ncSSc patients from controls, but not lcSSc.

**Conclusion:** We are the first to show that transcriptomic analysis of classical and non-classical circulating monocytes can unbiasedly stratify SSc patients and correlate with disease activity outcome measures.

## INTRODUCTION

Systemic sclerosis/scleroderma (SSc) is a complex, autoimmune-mediated connective tissue disease characterized by widespread vascular damage, chronic inflammation, and fibrosis. SSc is highly heterogenous in presentation, and may involve the skin, blood vessels, lungs, heart, kidneys, and gastrointestinal (GI) tract. There are two subtypes of SSc, limited and diffuse, with limited SSc representing 80% of all cases and diffuse SSc representing 20%. While there are no known cures for SSc, many studies suggest that mycophenolate mofetil (MMF/Cellcept), an immune modulatory drug that attenuates proliferation (1), is beneficial for SSc skin and lung disease (2–11). However, treatment failure or intolerance to MMF is common. Nonetheless, to date, there are no validated biomarkers to guide the selection of therapy to prevent end-organ disease in SSc patients, which represent a major unmet need for SSc patients. While clinically defined SSc subsets are associated with internal organ complications and death, they cannot predict disease course on a per-patient basis or inform treatment decisions. Thus, identification of disease endophenotypes and their longitudinal course will facilitate precision medicine implementation.

The role of immune cells in SSc pathogenesis is supported by the fact that the vast majority of genetic susceptibility loci in SSc belong to immunological pathways (12). Inflammatory cell infiltrates are observed in the skin by histology before fibrosis occurs (13), and changes to leukocyte frequency and function in the blood are accompanied by an increase in pro-inflammatory molecules including IL-6 (14–16). Additionally, fibrotic features in SSc patients reverse or stabilize after immunoablation followed by immune reconstitution with autologous CD34+ stem cells(12, 17–19). Circulating monocytes exist in 3 main states, characterized by CD14 and CD16 in humans: classical (CM) (CD14^++^CD16^-^), intermediate (IM) (CD14^+^CD16^+^), and non-classical (NCM) (CD14^dim^CD16^+^) (20, 21). The numbers of CD14^+^ monocytes in blood of patients with SSc are higher in SSc patients compared to healthy controls (22, 23) and are associated with reduced survival in SSc (23). Flow concentrations of circulating pro-fibrotic monocytes, especially the CD14^+^CD163^+^CD204^+^ cells (24), and fibrocytes (likely immature macrophages) are also elevated in patients with SSc-related interstitial lung disease (SSc-ILD) compared to healthy controls (24). Our group was the first to identify a causal role for monocyte derived alveolar macrophages in a murine model of fibrosis and to confirm the presence of these cells in humans with SSc-induced lung fibrosis (25–27). Increased numbers of dermal macrophages have been identified in skin from SSc patients (23) and in murine models of SSc-like disease (28). The numbers of CD163^+^ and CD204^+^ skin macrophages are higher in SSc patients as compared to healthy controls (24). Moreover, reductions in skin macrophage numbers are detected in a subset of SSc patients on MMF (29). These data suggest that monocyte/macrophage population is crucial for SSc development. However, little is known regarding individual populations of monocytes in SSc.

Patients with SSc display a complex clinical phenotype. Over the past several years, GWAS studies uncovered numerous susceptibility loci that contribute to the development of SSc. A genomic risk score using the GWAS data is sufficient to identify patients with SSc vs controls or other immune-mediated inflammatory disease (30). However, the GRS does not predict which individuals will develop SSc disease or the subclassification of SSc, most likely due to the high environmental contribution to pathogenesis. There are numerous studies that relate transcriptional signatures from PBMC or whole skin of SSc patients to disease activity (23, 31). We identified four molecular pathway-centric ‘intrinsic SSc subsets’ (inflammatory, fibroproliferative, limited and normal-like) using whole-genome microarray analysis of PB, whole skin and esophageal biopsies from SSc patients (32–35), yet only the inflammatory group in skin has been independently validated (36). We show that patients with an inflammatory gene expression signature in skin are the most likely to demonstrate skin disease improvement during MMF (37). However, there were inflammatory groups of patients who were non-responders (37, 38), which suggests that an understanding of the transcriptional profile of bulk immune populations may be necessary to discern which patients are more susceptible to a particular therapy.

In this study, we profiled the transcriptional signature of CM and NCM in SSc compared with age-, sex-, and ethnicity-matched controls within the Prospective Registry of Early Systemic Sclerosis (PRESS) consortium (39–44). Based on RNA sequencing (RNA-seq) data from CM, we define in an unbiased manner three sub-groups (A-C) of patients who may be clearly delineated by upregulation of distinct functional pathways including type 1 interferon, IL-1β, and TGF responsive clusters, respectively. While two of the sub-groups were recapitulated in NCM, monocyte maturation from CM to NCM phenotype differs in these three patient groups. The transcriptional clusters associated with groups A and C were also retained in bulk macrophages from skin of SSc patients. Group A correlated with highest forced vital capacity (FVC) at baseline, while group B had the lowest FVC even though there was no difference in mRSS in any group. Taken together, our study demonstrates the potential for circulating monocytes as a useful tool to stratify patients prior to therapeutic intervention.

## METHODS

### Study Participant Recruitment & Sample Collection

Patients in this study were recruited to the Prospective Registry of Early Systemic Sclerosis (PRESS) study at Northwestern University (NU), University of Texas, University of Utah, and University of Michigan (STU00062447). PRESS patients have early (defined as less than 2 years since the onset of the first non-Raynaud symptom attributed to SSc) diffuse cutaneous (dc) SSc (swollen/puffy hands or sclerodactyly and positive anti-RNA polymerase III or Scl70 antibodies and/or skin thickening involving the upper arms, thighs, or torso, and/or presence of tendon friction rubs, and absence of anticentromere antibodies). Clinical information on disease status was recorded for each patient. (Table 1). Age-, sex-, and ethnicity-matched controls were recruited at NU through the Control Blood Acquisition for Rheumatology Research (STU00045513) protocol. Whole blood from patients and controls was collected in EDTA tubes. For NU patients, blood was stored overnight at room temperature on a rocker and processed within 24 hours. Samples from remote sites were shipped overnight at room temperature and processed within 24 hours from the original shipped time. For skin biopsies, patients and healthy controls were recruited independently from NU (STU00002669; Table 1). Two side-by-side 4 mm dermal punch biopsies from the dorsal surface of the forearm midway between the ulnar styloid and the olecranon were sampled. Biopsies were immediately placed in MACS Tissue Storage Solution (Miltenyi Biotec) on ice.

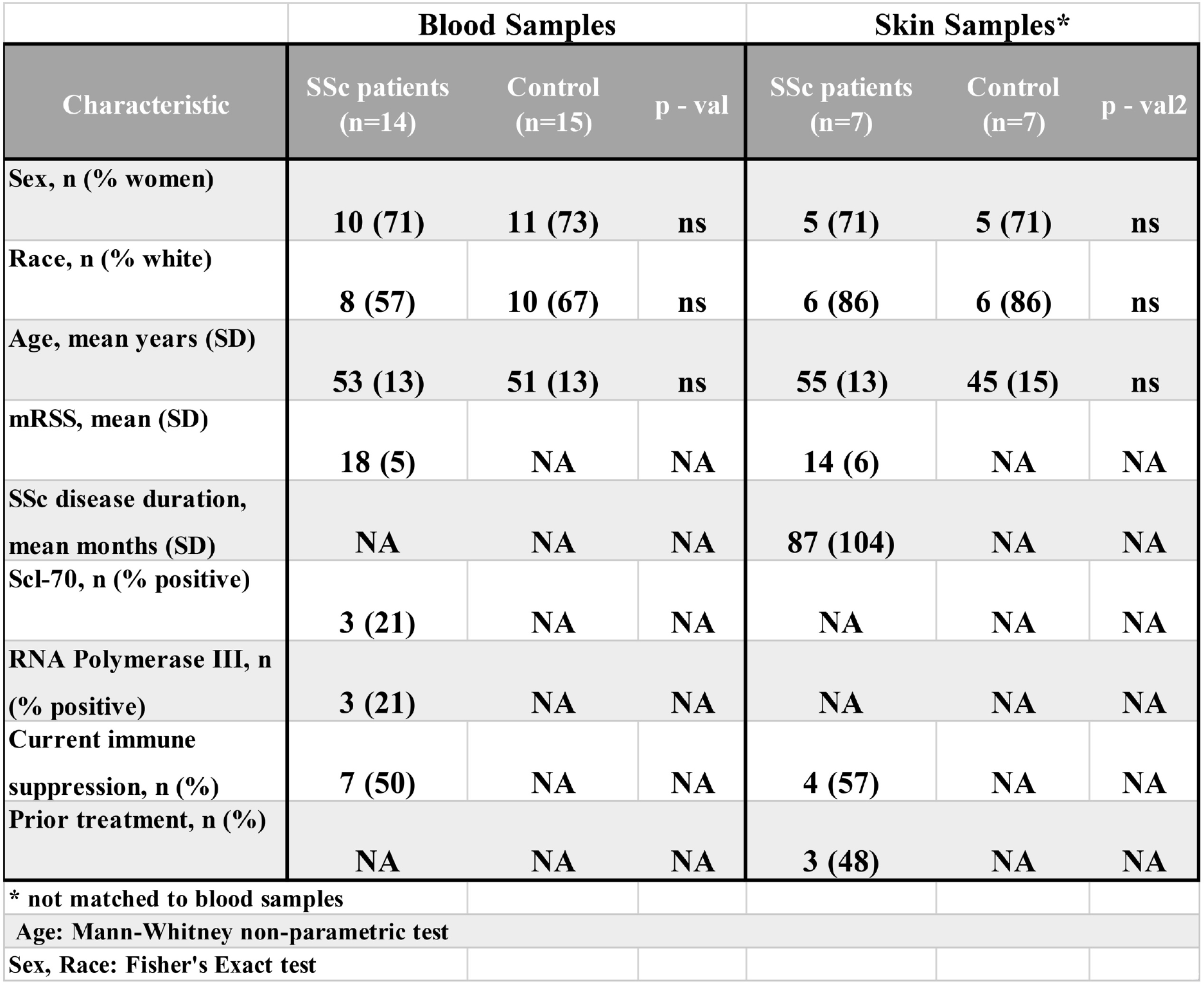

### Processing and Fluorescence Activated Cell Sorting (FACS) of Blood

Blood and tissue were processed, stained, and sorted via FACS as previously described (45). Twenty milliliters (mL) of whole blood were incubated for 8 minutes with gentle shaking in 80 mL red blood cell (RBC) lysis buffer (155 mM NH4Cl, 10 mM KHCO3, 0.1 mM EDTA in deionized water; warmed to 37 degrees). The sample was centrifuged at 350xg for 10 minutes at room temperature and RBC lysis was repeated on the pelleted cells. Lysis was quenched with MACS buffer (Miltenyi Biotec). Cells were washed with HBSS and counted (Invitrogen Countess). Ten million cells were stained with Fixable Viability Dye (0.5 uL/mL eBioscience) for 15 minutes in the dark at room temperature, then washed twice with magnetic-activated cell sorter (MACS) buffer (Miltenyi Biotec). Cells were incubated with FcR Blocking Reagent (Miltenyi Biotec; 1 *μ*L in 50 *μ*L MACS) for 20 minutes and then with a blood antibody cocktail for 30 minutes in the dark at 4 degrees: anti-CD45 BB515 (BD Horizon), anti-CD14 PerCP-Cyanine5.5 (eBioscience), anti-HLA-DR eFlour 450 (eBioscience),anti-CD15 APC (Biolegend), anti-CD16 APC-Cy7 (BD Biosciences), anti-CD1c PE (Miltenyi Biotec), anti-CD20 Alexa Flour 700 (BD Biosciences), anti-CD56 PE-Cy7 (BD Biosciences). Cells were washed twice and re-suspended in 1 mL MACS buffer and stored on ice. Cell sorting was performed on the FACSAria III (BD Bioscience) at Northwestern University RLHCCC Flow Cytometry Core Facility using a 100 μm nozzle, flow rate of 1, and 40 psi pressure. Populations were sorted into 100 μL MACS buffer, pelleted, re-suspended in 100 μL PicoPure RNA extraction buffer (Arcturus Biosciences), and stored at -80°C until RNA extraction.

### Skin Biopsy Digestion and Macrophage Sorting

Fat was discarded, and epidermal and dermal tissues were separated, infused with digestion buffer (RPMI 1640 [Sigma], Liberase TL [0.1 mg/mL; Roche], and DNase [0.1 mg/mL; Roche]) and minced with scissors. Tissue suspensions were transferred to C-tubes (Miltenyi Biotec) and incubated for 2 hours at 37°C. Tissue was mechanically digested with aggressive disaggregation using the pre-set gentleMACS program m_lung_01 before and after incubation. The digestion reaction was quenched with MACS buffer (Miltenyi Biotec), and the tissue suspension was filtered over a 40-micron filter. Cells were counted and all extracted cells were labeled with viability dye, treated with Fc block as described above, and stained with the following cocktail for 30 minutes at 4°C in the dark to enable delineation of mature mononuclear phagocytes: Anti-CD45 PE-Cy7 (BD), anti-HLA-DR eFluor 450 (eBioscience), anti-CD206 APC (BD Biosciences), anti-CD16 PE (BD Biosciences), anti-CD56 Alexa 700 (Biolegend), and Anti-CD15 Alexa700 (BD Biosciences). Cells were kept on ice until sorting as described above. Gated myeloid cells were sorted directly into 100 μL PicoPure and stored at -80°C until RNA extraction.

### Preparation of RNA Library

RNA from sorted cells was extracted using the PicoPure RNA isolation kit per manufacturer’s instructions (Arcturus). RNA quality and quantity were measured using a High-Sensitivity RNA ScreenTape System (Agilent Technologies). Library construction and RNA sequencing was performed in the Division of Rheumatology and Pulmonary and Critical Care sequencing facility. Full-length RNA-seq libraries from CM and skin macrophages were prepared from ∼3 ng of total RNA using SMART-Seq v4 Ultra Low Input RNA Kit (Clontech Laboratories) followed by Nextera XT protocol (Illumina). For NCM samples, the QuantSeq FWD (Lexogen GmbH, Vienna, Austria) was used to generate Illumina compatible libraries. Both full-length and 3’ RNA-seq libraries were sequenced on a NextSeq 500 instrument (Illumina) with 5-10 x 10^6^ aligned reads per sample. A commercially available universal human RNA (uhRNA) reference was prepared along with sample RNA to represent background expression (45).

### RNA seq Analysis

Sequencing data were de-multiplexed using bcl2fastq and reads were aligned to the human reference genome (NCIB, hg19) using TopHat2 (version 2.17.1.14) for full-length data sets (CM and skin myeloid cells) or STAR (version 2.5.2) for 3’ data sets (NCM) (46, 47). Aligned reads were mapped to genes and counted using HTseq (48). Gene coverage (5’à3’) for full-length libraries were calculated using the RSeQC package (49). Gene counts were normalized by calculating fragments per kilobase per million (FPKM) for CM and skin macrophage samples or counts per million (CPM) for NCM samples. In CM and skin samples, expressed genes were defined as genes for which log2(FPKM+1) ≥ 3 in at least three samples. NCM expressed genes were also defined as log2(CPM+1) ≥ 5 in at least 3 samples. Samples with fewer than 5 x 10^6^ singly mapped reads or with complexity (percentage of reads mapping to unique genomic positions) less than 30% were excluded.

### Computational Analysis and Visualization

R Studio (https://www.rstudiocom/) was used to calculate Pearson’s correlation coefficients and to generate PCA plots, Venn diagrams (ggplots package), and scatterplots (ggplot2 package). PRISM (GraphPad Software LLC) was used to generate all bar graphs and line plots. Differential expression was calculated using the DEseq package (version 1.18.1); genes with changes in expression of two-fold or greater and Benjamini Hochberg adjusted p-value < 0.05 were considered differentially expressed genes (DEGs) (50). To define genes differentially expressed in a subset of SSc patients compared with healthy controls in CM, we calculated a modified z-score (*z*′) for each gene based on its distribution of expression in the control cohort:

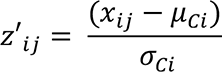

Where *x* is the expression level of gene *i* in patient *j, μ*_*ci*_ is the mean expression for gene *i* across controls, and σ_*Ci*_ is the standard deviation of expression for gene *i* across controls. To prevent artificially high z scores resulting from small standard deviations, where high z score may not represent true biological variability, we set the minimum σ_*Ci*_ to 2.5. We identified 1790 genes with |*z*′| > 2 in at least 3 patients. GENE-E (https://software.broadinstitute.org/GENE-E/) was used to perform hierarchical clustering of samples and K-means clustering of genes (k=4), resulting in patient groups A-C and gene clusters I-IV. Pvclust (51) was used to assess the uncertainty in hierarchical clustering analysis. For each cluster in the hierarchical clustering, p-values were calculated via multiscale bootstrap resampling. P-value of a cluster (between 0 and 1) indicates how strong the cluster is supported by data and the AU (Approximately Unbiased) p-value in red and BP (Bootstrap Probability) value in green are reported. P-values for the SSc patients (CM and NCM) shows how confidently the clusters are supported by the underlying data when clustered hierarchically.

To evaluate the robustness of the three hierarchical groups of patients and determine if the classification held up or changed if a randomly sampled subset of our dataset was used, we employed a bootstrapping approach of analysis. We used varying subset sizes ranging from 5% to 95% of the entire dataset (out of 5171 genes) to re-cluster the patients using hierarchical clustering. For each of the subsets selected randomly for 1000 iterations, we computed the success rate (by counting how many times out of 1000 were the same patients clustered together for each of the 3 groups) for obtaining the same 3 clusters (same groups of patients classified together) that we identified for the full dataset. We then plotted the success rate (%) for obtaining each of the individual groups correctly with the varying subset sizes as well as the success rate for obtaining all 3 groups correctly. Through this bootstrapped evaluation approach, we verified the extent to which randomized chunks of the full dataset can reproduce the 3 groups. We also verified that our groups do not form hierarchical clustering by chance but rather due to the underlying biological variance within the patients in those groups.

### GSEA Analysis

To determine the biological processes of enriched genes in each group, genes were ranked by group based on their fold-change relative to the mean expression across controls and given as pre-ranked input for Gene Set Enrichment Analysis (GSEA v3.0, www.gsea-msigdb.org). Pre-ranked genes were compared to the biological processes (BP) of the Molecular Signature Database (MSigDB) C5v7.0 collection using pre-ranking and classic weighting. Normalized enrichment scores (NES) were reported, and significant enrichment (or depletion) was defined as adjusted p value < 0.05.

### NCM analysis

We calculated the modified z-score of the identified CM, in NCM and identified 1637 of which were also expressed in NCM. Hierarchical clustering was performed on NCMs from patient’s samples based on these scores. Since groups were maintained, correlation analysis of genes between CM and NCM were performed, cell specific conserved genes were delineated from genes that were conserved across both cell subsets. Enriched Gene Ontology (GO) terms of genes that were conserved across both cell subsets were determined using GOrilla (cbl- gorilla.cs.technion.ac.il) (52) and plotted using ggplot2.

### Canonical monocyte gene comparison

We performed a comparative overlap analysis of our CM and NCM to publicly available canonical gene data(53). We identified 431 CM genes and 388 NCM genes. We then analyzed the percentage of genes that more closely represented in the canonical genes per group, compared to control subjects. Data were represented with bar graphs and average expression of canonical genes per group were represented with dot plots.

### Skin Macrophages

We used a non-parametric Mann-Whitney U test to identify significantly up-regulated (190 genes) and down-regulated (138 genes) genes in skin macrophages from patients compared to controls and FPKM values were plotted in heatmap. Representative gene expression was shown using bar graphs. Number of skin macrophage genes that overlap with CM genes were calculated in patients and controls and shown using bar graphs. ROC curve, (pROC package)(54) was used to determine the efficacy of gene Clusters I-IV at delineating patient vs control status. Samples were given a cumulative score as the sum of the min-max value for all genes in the set, p-values are calculated by the package using the Mann-Whitney U statistics addressing the null hypothesis $H_o:$(55).

### Statistical analysis of clinical data

Baseline FVC and mRSS differences among the groups identified in the cluster analysis were assessed using the Kruskal-Wallis test. FVC and mRSS differences at baseline between Groups A and the combination of Groups B and C were assessed using a Mann-Whitney U test. Baseline distributions of FVC and mRSS by groups identified in the cluster analysis were illustrated using box plots. Additionally, the longitudinal FVC and mRSS data are displayed in spaghetti plots.

### Comparison with published SSc data

CM cluster I-IV were compared to published data set of whole monocytes obtained from SSc patients. All information on patient diagnosis, grouping and sample acquisition are available publicly by van der Kroef et al (56). Overlapping genes between the public SSc data and our clusters were then given a cumulative score as the sum of the min-max value for all genes per sample per cluster, then the SSc patients data were plotted for each cluster. ROC curves were plotted to ascertain the delineated clusters of the overlap analysis.

## Results

### Classical Monocytes from SSc Patients Exhibit Greater Variability than Healthy Controls

We obtained blood samples from 15 SSc patients and 15 age, sex, and race-matched healthy volunteers used as controls (**Table 1**). We then isolated classical (CM) and non-classical (NCM) monocytes and found that there were no significant differences in the proportion detected among CD45^+^ immune cells between the SSc and control cohorts (**Figure 1A-B**). We performed RNA-seq to profile the gene expression of CM, the predominant monocyte population, eliminated one patient sample that did not pass quality control and confirmed the gender of the patients (**Supp. Figure 1A-F**). From the remaining samples, we found that the expression of genes tended to be more variable across SSc patients than across controls (**Figure 1C, Supp Figure 1C**). By calculating an average transcriptional profile for the control cohort, we determined that SSc patients exhibited a range in their similarity to “healthy control” (**Figure 1D**). Moreover, while the patients that most resembled controls were necessarily similar to each other, the remaining patients exhibited substantial variability in their transcriptional profiles (**Figure 1E, Supp Figure 1E**). This variability did not seem to be associated with the site of sample collection, although patients currently being treated with immunosuppressive therapies tended to more closely resemble controls (**Figure 1F**, **Supp Figure 1F**). We next performed pairwise differential expression analysis between the SSc and control cohorts and defined 263 and 66 genes as up- and down-regulated in SSc, respectively (**Figure 1G**). The 263 included genes involved in type I interferon signaling pathway (*OAS1* and *IRF2*), cAMP signaling (*FOS*, *FOSB* and *JUN*), regulation of angiogenesis (*VEGF* and *CXCR4*) and regulation of chemotaxis (*CSF1R*, *CCR2* and *CCL2*). Although these results agree with prior reports (57), the high variability in the SSc cohort suggests that a pairwise approach does not accurately reflect all patients.

**Figure 1:**
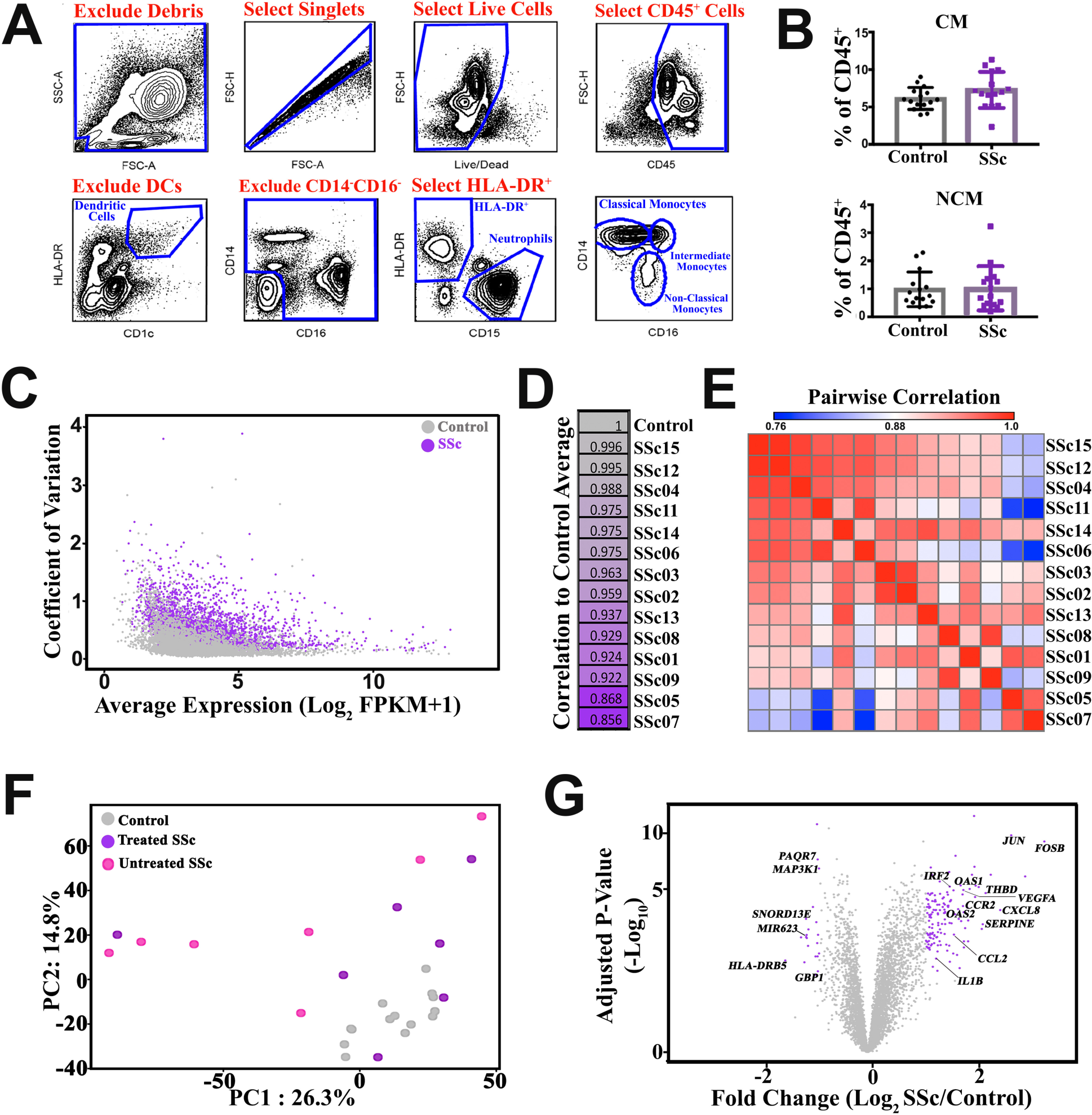
Classical Monocytes from early diffuse SSc patients display transcriptional heterogeneity. **(A)** Contour plots depicting the step-by-step isolation of classical monocytes (CM) and non-classical monocytes (NCM) from blood by FACS. **(B)** The percent of CM and NCM in CD45+ cells from SSc compared to control samples (p = 0.14 and 0.89, respectively. Mann- Whitney test). **(C)** Scatterplot showing the coefficient of variation (standard deviation/average FPKM) relative to the average expression (log2(FPKM+1)) in SSc (purple) and control (grey) CM. **(D)** Correlation (Pearson’s coefficient) of CM expression in each SSc patient relative to the average gene expression across control samples. **(E)** Pairwise Pearson correlation between gene expression of CM samples from SSc patients. **(F)** Principal Components Analysis (PCA) of gene expression in CM samples from Treated patients (purple), Untreated patients (pink), and controls (grey). **(G)** Volcano plot showing differentially expressed genes in the SSc cohort compared with controls log2 fold change >= 1 or <=-1 (significance calculated by DEseq with Benjamini Hochberg FDR correction, purple indicates genes with Padj < 0.05 & log2 fold change >= 1 or <=- 1). **C-G** based on 5171 expressed genes in CM.

### SSc Classical Monocytes Cluster into Three Transcriptional Subgroups

In order to capture the transcriptional dysregulation in individual SSc patients, we devised an alternative approach to define differentially expressed genes. For each patient, we considered a given gene to be differential if its expression was more than 2 standard deviations from the control mean (i.e. adapted z-score > 2 or < -2; see Methods). By hierarchical clustering of the SSc patients based on the 1790 genes that were differential in at least 3 patients, we identified 3 patient subgroups (A-C) excluding 1 outlier patient (SSc15) that was highly similar to controls (**Figure 2A**). The clustering of patients was robust: while individual branches of the dendrogram changed, the identity of the groups remained largely constant regardless of the subset of genes used (**Supp Figure 2A-B)**. Moreover, by k-means clustering of the 1790 genes, we characterized 4 expression patterns associated with these subgroups: Cluster I is primarily expressed in Group A, Cluster II in Group B, Cluster III in Group C, and Cluster IV is down-regulated in the majority of patients compared with Controls (**Figure 2A-D**). The distinct transcriptional identities of each group were reinforced when visualized on the PCA from **Figure 1** (**Figure 2E**). Patients in Group A and B tended to have the most differentially expressed genes (**Figure 2F**). On the other hand, Group C was associated with genes not detected previously and represent dysregulated pathways that may have been missed in earlier studies as well.

**Figure 2:**
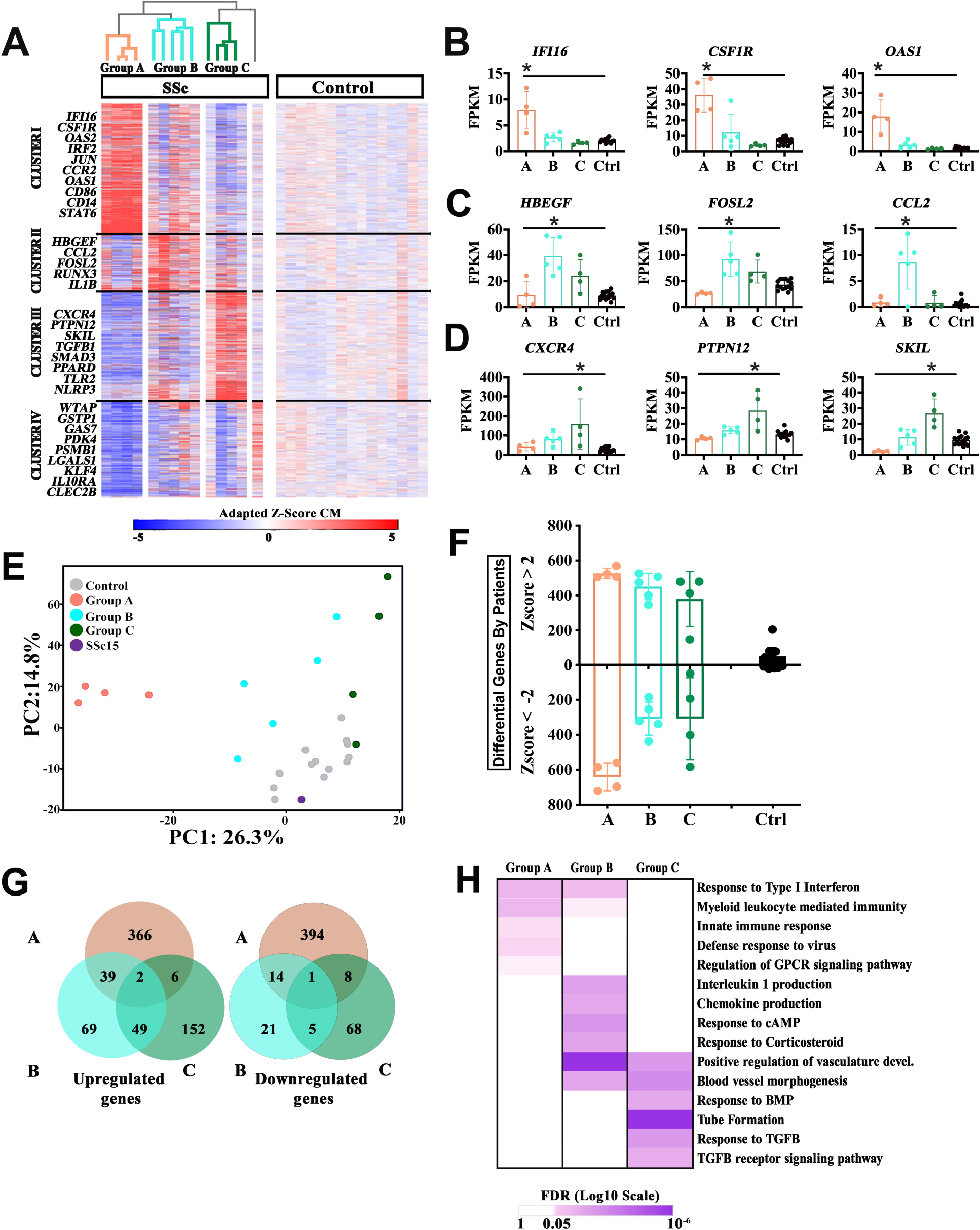
Early diffuse SSc patients stratify into 3 groups based on the transcriptional profile of CM. **(A)** Heatmap of adapted z-score of CM gene expression relative to controls, depicting dendrogram resulting from unsupervised hierarchical clustering of samples (columns) into 3 groups (Group A=coral; Group B=cyan; Group C=green) and K-means clustering of 1790 differential genes by individual (|adapted z-score| > 2 in at least 3 patients) (rows) (K=4). Expression of representative genes from **(B)** Cluster I (592), **(C)** Cluster II (261), **(D)** Cluster III (500), and **I** Cluster IV (437) in each patient group as well as controls. * Indicates padj < 0.05 calculated by DEseq with Benjamini Hochberg FDR correction **(F)** Number of genes with adapted Z scores greater than 2 or less than -2 in individual CM samples organized by patient groups. **(G)** Venn Diagram showing overlap of upregulated ((log_2_FC >1, and p-adj < 0.05) or downregulated ((log_2_FC <-1, and p-adj < 0.05) genes in each patient group compared to controls. **(H)**Heatmap showing the False Discovery Rate (FDR) q-values of biological processes associated with differential genes in the three patient groups (GSEA). **(I)** PCA of gene expression in CM samples color-coded based on patient groups. **A-I** based on 5171 expressed genes in CM.

We performed pairwise differential analysis on each of the patient groups defined by hierarchical clustering to better characterize their transcriptional profiles (**Figure 2G-H, Supp Figure 2C, E, G**). As above, we found that Group A exhibited the most differentially expressed genes and were associated with high levels of type I interferon responsive genes – such as *IFI16*, *OAS1*, *OAS2*, and *IRF2 –* as well as innate immunity in general and GPCR signaling including *CSFR1*, *CCR2* and *CD86*. Group B had the fewest differentially expressed genes, the most overlap with other groups. This may also explain why it was the least conserved in re-sampling (**Supp Figure 2B**). Like Group A, Group B genes were associated with type I interferon pathway and innate immunity, but exhibited higher enrichment of chemokine production and response to cAMP and corticosteroids, including *CCL2*, *IL1B*, *RUNX3* and *FOSL2*. Both Group B and C upregulated genes associated with vasculature development and blood vessel morphogenesis, such as *VEGFA* and *HBEGF* (58), but Group C additional genes associated with angiogenesis (i.e. *CXCR4* (59) and *PTPN12* (60)). Further, Group C also expressed upregulated genes associated with TGFB and BMP signaling, including *TGFB1*, *SMAD3*, *SKIL* and *PPARD*. Taken together, these results demonstrate that CM from SSc patients can be classified into subtypes with distinct transcriptional identities.

### Patient Subgroups are Largely Conserved in Non-Classical Monocytes

Next, we performed RNA-seq on NCM isolated from the same patients (**Supp Figure 3A**). The variation in gene expression was higher in NCM than CM in both patients and controls (**Supp Figure 3B-D**), but this may be due to technical differences in the protocols or higher sampling bias from the smaller NCM population (< 5% of total monocytes). When NCM were clustered based on the same 1790 genes as CM, we found that the gene expression signature distinguishing patient groups was still evident (**Figure 3A**). Patients in Group A and C still robustly cluster together using these genes, but Group B no longer forms its own clade (**Supp Figure 3E-G**). As with CM, we performed pairwise differential expression on each group (**Supp Figure 3H**). We then compared the differentially expressed genes in both monocyte populations for each group (**Figure 3B-D**). We found that genes related to interferon and innate immunity were preferentially expressed in Group A in both CM and NCM (Figure 3B, E, and F and Supp Figure 3I). Although Group B did not exhibit as many genes overlapping between CM and NCM (i.e. *CCR2*), those that overlapped were associated with similar processes as CM alone (Figure 3C, E, and F and Supp Figure 3I) . Finally, Group C exhibited far fewer differential genes in NCM: in particular, genes related to TGFB signaling were not upregulated in Group C NCM (Figure 3D, E, and F and Supp Figure 3I). These results confirm that the transcriptional identity of patient groups extend beyond CM; however, differences between monocyte populations in SSc may be evidence of cell-type specific roles in disease.

**Figure 3:**
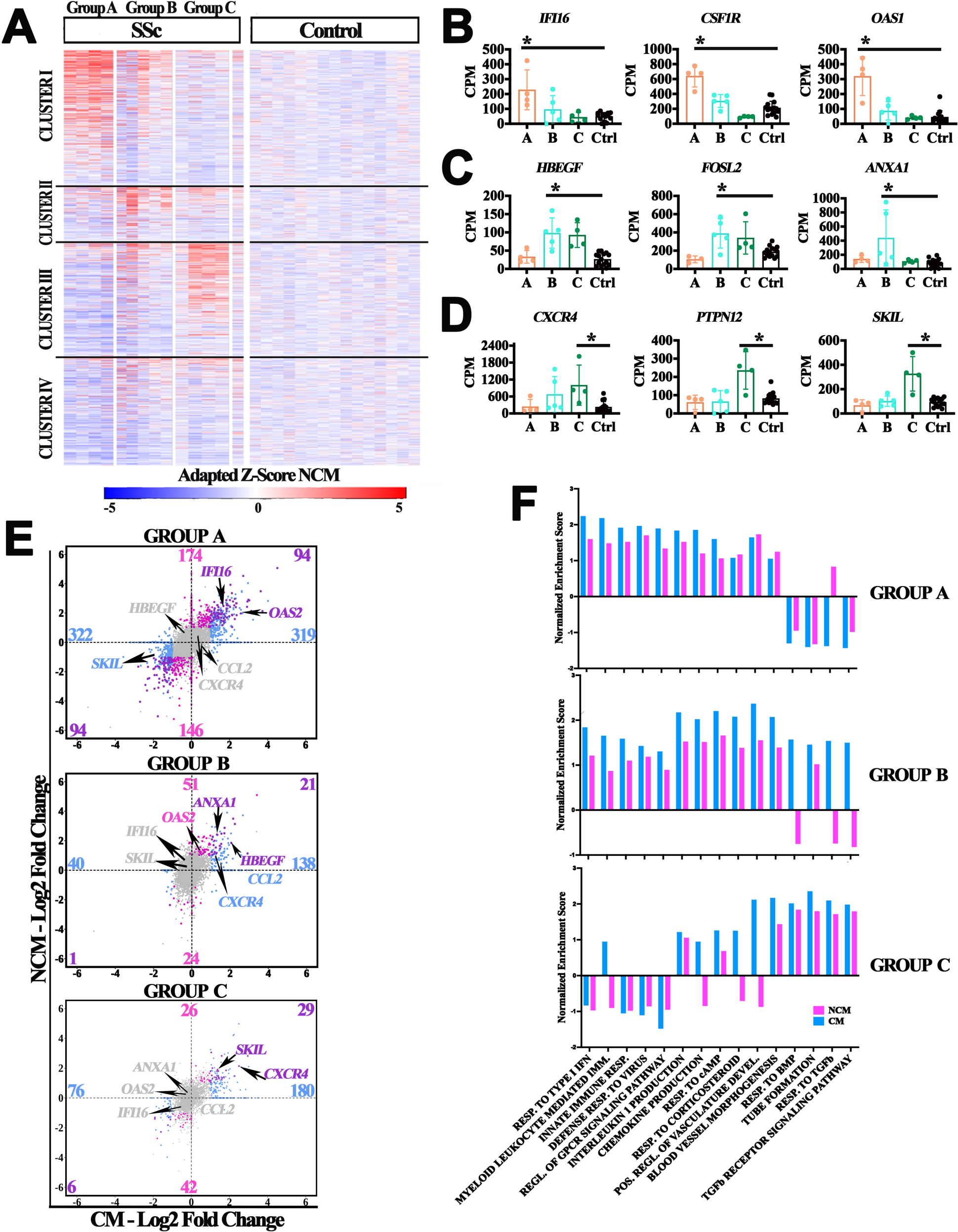
Patient groups from CM exhibit distinct transcriptional profiles in NCM. **(A)** Heatmap of adapted z-score of NCM gene expression relative to controls using 1599 genes from CM clusters in 2A. **(B-D)** Expression of representative genes from clusters I-III respectively. * Indicates padj < 0.05 calculated by DEseq with Benjamini Hochberg FDR correction **(E)** Scatterplot of Log_2_FC of gene expression in each patient group compared to controls in CM (x-axis) versus NCM (y-axis). Significant genes (Log2FC =>1 or <= -1and Padj<=0.05) are colored if they are shared between CM and NCM (purple), only NCM (pink), or only in CM (blue). **(F)** Bubble plot showing the biological processes associated with the genes that are upregulated across both cell types**. A-F** based on 7143 expressed genes in NCM.

### Monocyte identity is Altered Across Patients

It has been proposed using mouse models that NCM differentiate from CM that have egressed from the bone marrow (61). The genes that distinguish NCM from CM on a transcriptional level have been well-established and shown to be conserved between mouse and humans (62). To determine whether monocyte identity is altered in disease, we generated a list of CM-specific and NCM-specific genes based on recent studies (63, 64). We found that CM from Groups A and B patients expressed canonical genes, including *PLAC8*, *PADI4*, *SELL*, *LYZ*, and *CCR2*, at a higher level on average than controls (**Figure 4A, C**, **Supp Figure 4A, C**). Group A NCM also expressed higher than average levels of their canonical genes, including *ITGAL*, *SPN*, *CX3CR1*, and *CSF1R*, than controls (**Figure 4B, D**, **Supp Figure 4B, D**). In comparing CM and NCM, we may have expected that the genome-wide transcriptional profile would be most similar between monocytes from the same patients; however, this is not the case (**Figure 4E**). The over-expression of CM genes in Groups A and B may explain why CM from these patients are less similar overall to NCM **(Supp Figure 4E, F)**. It is possible that these results stem from the disease pathways that are differentially activated in patients and affect monocyte differentiation.

**Figure 4:**
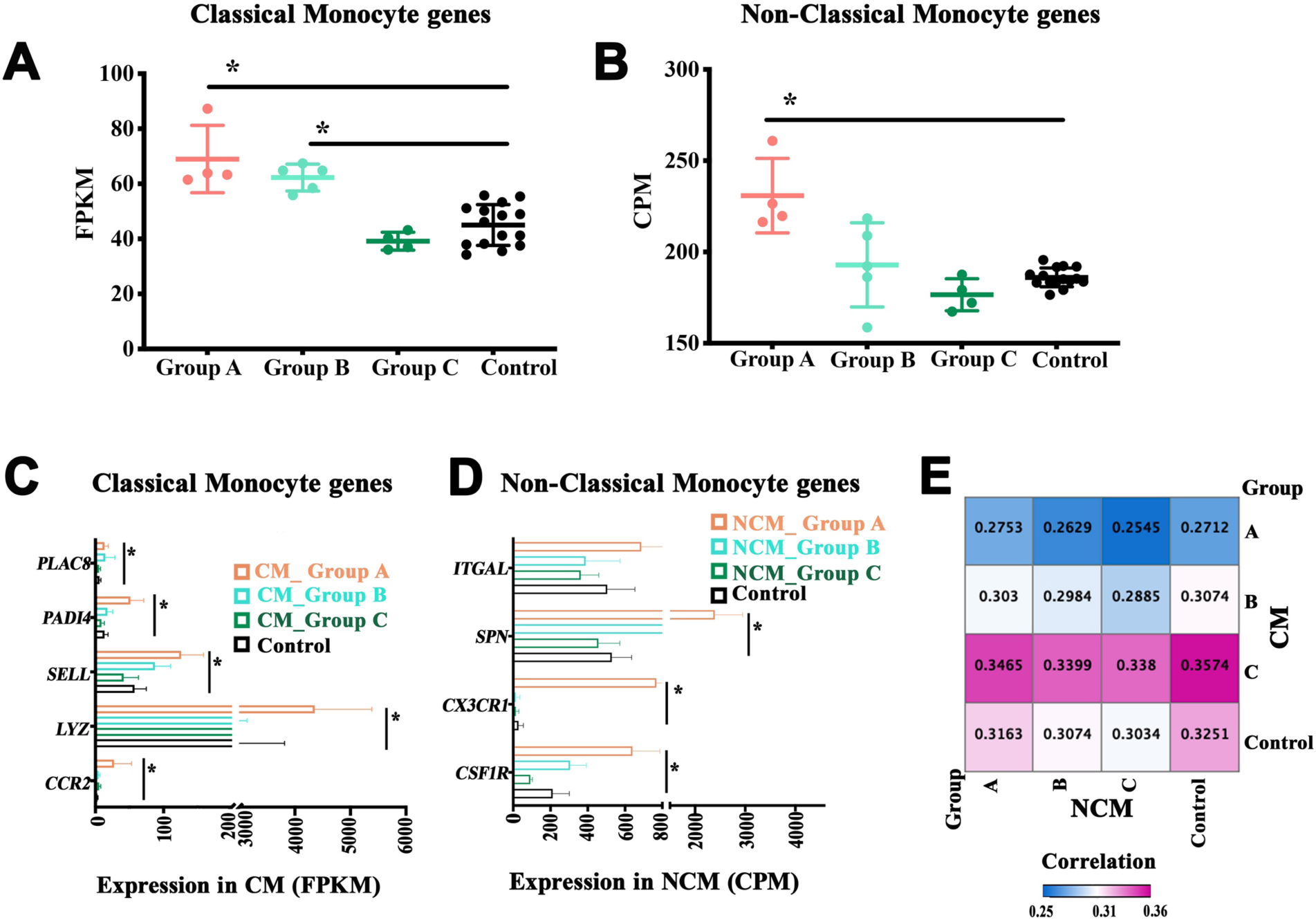
Variation in expression of monocyte gene signature in SSc CM and NCM. **(A)** Average expression level of all CM genes in CM samples of patient groups and controls (B) Average expression level of all NCM genes in NCM samples of patient groups and controls. * Indicates p < 0.5 Mann-Whitney nonparametric test. **(C)** Expression levels of representative CM genes in patient CM samples by group. **(D)** Expression levels of representative NCM genes in patient NCM samples by group. **(E)** Pearson correlation of average gene expression between CM and NCM samples in patient groups and controls.

### Skin Macrophages Upregulate SSc-Associated Monocyte Genes

Since we observed common transcriptional signatures across monocyte populations, we then investigated whether similar gene expression profiles could be observed in tissue macrophages. Monocytes and macrophages share many genes in common and prior studies have shown that SSc-associated fibrosis is driven by monocyte-derived macrophages in the tissue (65). We isolated CD206^+^HLADR^+^ macrophages from skin biopsies of SSc patients with evidence of fibrosis (**Figure 5A**). We performed RNA-seq on these cells (**Supp Figure 5A, B**) and identified 328 differentially expressed genes between skin macrophages in SSc patients vs. controls (**Figure 5B and Supp Figure 5C**). The 190 SSc-specific genes included those that play a role in myeloid leukocyte mediated immunity. More specifically, genes involved in cell adhesion, phagocytosis, or innate immune response such as *ITGB2*, *ITGA5* and *ADAM8* (66), *ALOX15* (67) and *CLEC10* (68), and *NFKB2*, *NFKBIE* (69), *PIM1* (70) and *RUNX3* respectively. The 138 control-specific genes included transcription factors such *CEBPD*, which potentiates macrophage inflammatory response and angiogenesis (71, 72), *ID3*, which is necessary for macrophage specification in liver (73), *EPAS1* (HIF2a), which suppresses the *NLRP3* inflammasome (74) and *LRG1* (75) and *IL6*, which play a role in angiogenesis (**Figure 5C**). Of note, *ITGB2*, *ITGA5*, *ADAM8*, *CEBPD*, and *NFKB2* were all differentially expressed in CM SSc patients compared to controls. We found that Clusters I-III genes, which corresponded to upregulated genes in Groups A, B, and C in our monocyte analysis, were more likely to be SSc-specific than control-specific (Enrichment: I=1.54, II=1.82, III=3.27), (**Figure 5D, Supp Figure 5D**). Similarly, classifiers based on the average expression of Clusters I and III genes in skin macrophages were able to differentiate between SSc patients and controls significantly better than random (p= 0.03, 0.01) (**Figure 5E**). On the other hand, Cluster IV genes, which were largely downregulated in monocytes, were generally expressed higher in controls than patients (Enrichment: IV=0.40). Taken together, these results suggest that the transcriptional profile of monocytes in SSc patients is conserved in skin macrophages.

**Figure 5:**
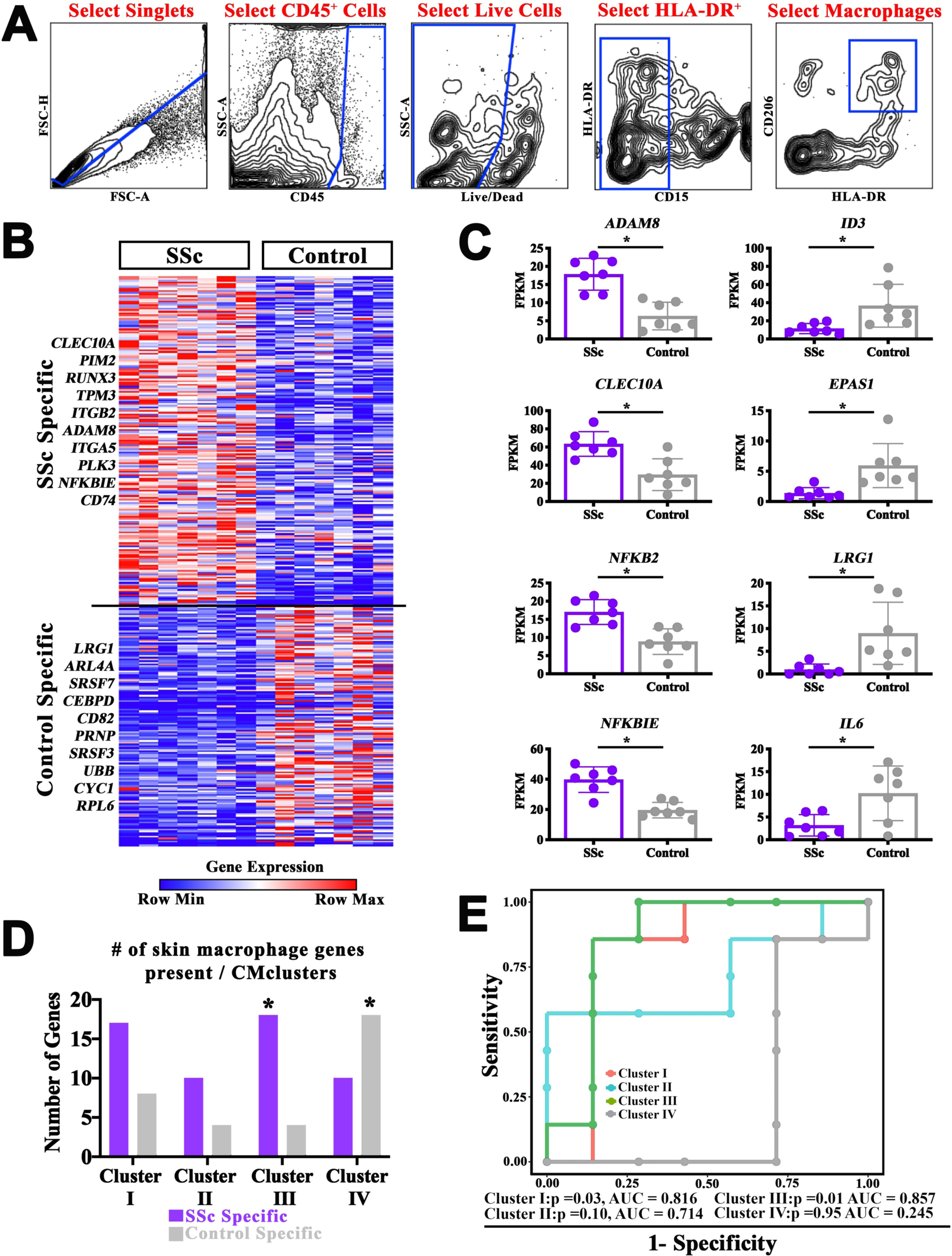
Clusters I and III genes are upregulated in SSc skin macrophages compared to control. **(A)** Gating scheme depicting the purification of macrophages from skin. **(B)** Heatmap of 190 SSc-specific and 138 Control-specific genes based on significantly differential expression in skin macrophages (p < 0.05 by Mann-Whitney U test). **(C)** Expression of representative genes from heatmap in skin macrophages from SSc patients and controls. **(D)** Number of genes from **B** that overlap CM Clusters I-IV in **2A**. **(E)** ROC curve based on the sensitivity and specificity of distinguishing skin macrophages from SSc patients vs. controls using the average normalized expression of genes from CM Clusters I-IV (**2A).** B-E based on 6338 expressed genes in skin macrophages.

### Patients Subgroups Differ in Disease Characteristics

Our analysis stratified SSc patients based on their transcriptional profiles in monocytes but was agnostic to their clinical presentation. We followed patients up to 42 months after the initial blood samples were obtained, and recorded their pulmonary function by Percent Forced Vital Capacity (FVC) and skin fibrosis by modified Rodnan Skin Score (mRSS). We found that Group B and C patients exhibited significantly worsened lung function at baseline than Group A patients and did not tend to improve over time (**Figure 6A, C**). In contrast, there were no significant differences in mRSS levels between groups at baseline or longitudinally (**Figure 6B, D**). These initial results indicate there may be phenotypic differences between our patient groups defined based on their transcriptional profile that can be verified in a larger longitudinal cohort.

**Figure 6:**
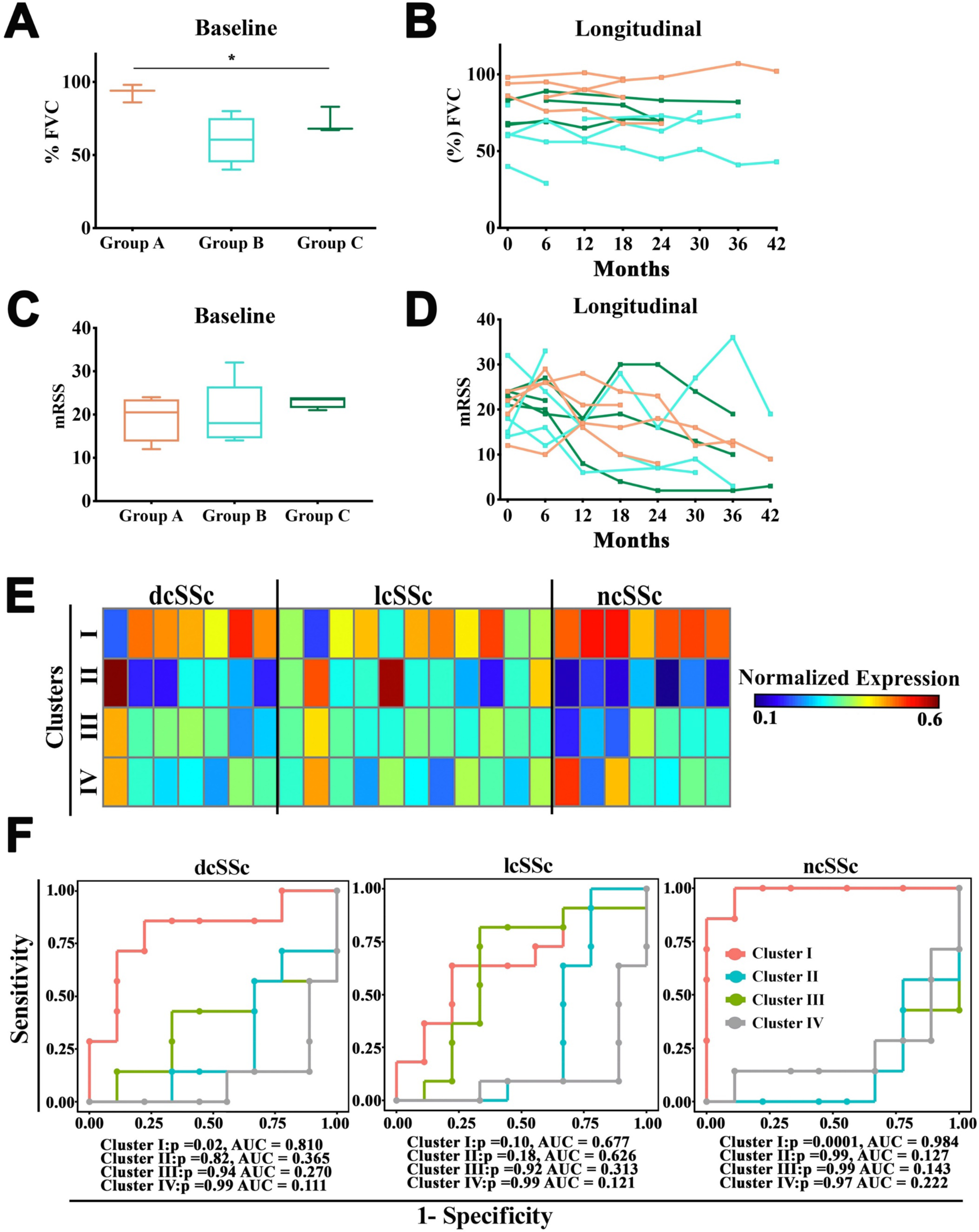
SSc patient groups differ across clinical phenotypes. **(A)** Forced vital capacity (FVC) and (**C)** modified Rodnan Skin Score (mRSS) at baseline (time of blood sample) of each patient by group, * Kruskal-Wallis test for 3 groups. **(B & D)** Longitudinal measurements of **(B)** FVC and **(D)** mRSS of each patient over 42 months from baseline (time 0). Patients are color-coded by group. **(E)** Average normalized expression of genes from CM Clusters I-IV (**2A**) in bulk monocytes from SSc patients categorized by disease phenotype (dataset described in van der Kroef, et al). **(F)** ROC curve based on the sensitivity and specificity of distinguishing bulk monocytes from SSc patient categories vs. controls using the average normalized expression of genes from CM Clusters I-IV (**2A)** (dataset described in van der Kroef, et al).

### Application of Transcriptional Groups to Independent Dataset

In order to apply our classification of patient groups to future studies, we aimed to develop an unbiased algorithm that is tolerant to differences in the underlying data resulting from technical artefacts, protocol choice, sequencing depth, and normalization. To test the algorithm, we used gene expression data from an unrelated study on monocytes in SSc patients performed by another group (56). In this study, they isolated bulk monocytes for RNA-seq, and categorized patients based on disease presentation: diffuse cutaneous (dcSSc), limited cutaneous (lcSSc), or non-cutaneous(ncSSc). For each patient, we calculated a score to reflect the relative level of expression of each cluster from our monocyte analysis (**Figure 6E, Supp Figure 6A**). We found that there was variability in the transcriptional signature of each patient that did not align with their disease category. We then built classifiers for each cluster and assessed their performance on patients categorized by disease presentation (**Figure 6F**). As expected, Cluster IV, which was downregulated in the majority of our patients, worked better as a negative classifier in each category. Cluster I (associated with Group A) accurately distinguished dcSSc from controls (p=0.02), but performed particularly well on ncSSc patients. Cluster I and II both exhibited high, but not significant AUC values in lcSSc, suggesting that this category is split between these transcriptional identities. Although individual patients exhibited high expression of Cluster III genes, it did not perform well as a classifier in any one category. These results suggest that the transcriptional signatures we defined can be identified in independent data sets, but more investigation is needed to determine the full clinical significance of our patient groups.

## DISCUSSION

Multiple studies have quantified gene expression in whole peripheral blood or skin from SSc patients (57). Like all bulk microarray or RNA-seq experiments, these studies are subject to changes in cellular composition that can drive gene expression signatures and a loss of the ability to detect biologically important transcriptional changes within minority cell populations. As such they rely on complex analyses, such as deconvolution and network analysis, and a priori knowledge from extant gene expression databases (e.g., GEO). Here, we are the first to show that unbiased transcriptomic analysis of CM and NCM can stratify SSc patients and correlate with disease activity outcome measures. One of the strengths of focusing on circulating monocytes is that they are composed of only 2-3 populations (76, 77), which allows for bulk separation. Patients segregated into three groups based on CM transcriptional profiles, with each group demonstrating upregulation of functionally distinct gene sets including response to type I interferon, myeloid leukocyte mediated immunity and regulation of GPCR signaling pathway for group A, Interleukin 1 and chemokine production, response to cAMP and response to corticosteroid for group B and tube/vessel formation, response to BMP, and TGF-ß receptor signaling pathway for group C compared to each other and to health participants. Since these three SSc endophenotypes are largely recapitulated in analysis of NCM transcription profiling, these data suggest some conservation is the gene signatures from Groups A, B and C. Analysis of bulk skin CD206^+^HLADR^+^ macrophages revealed that the majority of the genes associated with groups A, B and C are not present in skin macrophages (only 8 genes present); however, the central disease-associated GO pathways are similar between skin macrophages and circulating monocytes. These results suggest a connection between circulating precursors and mature tissue-resident cells, and ongoing studies are underway to replicate these results in paired skin and blood, which will enable direct comparison of immature and mature myeloid gene expression. The group A SSc patients, which expressed genes representing the response to type I interferon pathway, myeloid leukocyte mediated immunity and regulation to GPCR signaling pathway, display the highest FVC compared to either Group B or C but exhibits no change in mRSS. Together, our results suggest that transcriptional profiling of distinct CM and NCM subpopulations may represent a viable mechanism for identify patients and potentially their response to therapeutics.

Over the past decade numerous studies have utilized transcriptional profiling to understand SSc disease pathology or generate new patient stratification methods. Gene expression analysis of whole PB revealed a type I interferon signature as well as TLR and p53 pathways (31, 78). However, the type I interferon is also detected in whole blood from patients with SLE or polymyositis (78). DNA microarray of skin biopsies from patients with SSc has demonstrated four SSc “intrinsic” subsets that may be detectable in PBMCs and esophageal biopsies(34, 79, 80); however, it had not been established that this type of differentiation is detectable outside of diseased end-organs and to what extent immune cells drive this phenotype (29, 35, 37, 79, 81, 82). Further, an inflammatory gene signature associated with macrophages in bulk skin using RNA seq provides additional support for the crucial roles that macrophages play in the development of SSc (44, 83). Moreover, a change in skin severity score that includes a 415 gene signature has been shown to correlate with mRSS (84), although to date no studies have corroborated the skin severity score. Recently, a few groups examined gene expression specifically in individual immune populations such as monocytes or macrophages using bulk or single cell RNA-seq (56, 85, 86). While these studies support the role for the type I interferon as well as the IL-1β pathway in bulk SSc monocytes (CM and NCM), neither study examined any clinical associations (56, 85).

Further, the only single cell RNA seq study of human SSc skin examined the proportion of populations and their respective gene signature but did not perform any correlation analysis with disease outcome measures (86). The importance of our CM and NCM data are that it clearly illustrates the presence of disease subsets –or endotypes –in circulating immune cells. Further, our data is able to differentiate subtypes of SSc patients in another study, which are only stratified based their clinical classification i.e., dcSSc, lcSSc, and ncSSc (56). These data suggest that comparison of variability across populations which can be discerned via CM or NCM RNA seq analysis, may influence the traditional analyses and classification of SSc patients. Although our study clearly delineated three patient groups without apparent distinguishing clinical phenotypes, further studies will establish the reproducibility of this grouping and expand the sampling to definitively test any association with clinical parameters. Longitudinal analysis is needed to determine whether these groups are static with and without treatment.

Clinical heterogeneity in SSc presentation has been well characterized for decades, and differential gene expression has been demonstrated in fibrotic tissue; however, reproducible connections have not been established between transcriptomic intrinsic subsets and clinical characteristics, including prognosis and response to treatment. Approximately half of all organ manifestations in patients with SSc occur simultaneously within in the first two years following the onset of Raynaud’s phenomenon based on the longitudinal EUSTAR study (87). It is unknown whether the development of fibrosis in tissue is synchronous or asynchronous as few studies have assessed the evolution of organ involvement in SSc patients in a longitudinal manner. It has been proposed that a thorough molecular understanding of SSc would help to stratify patients for enrollment in clinical trials and to inform drug selection. Recently tocilizumab has been examined in SSc patients (88–93) as it is well known that circulating IL-6 is increased in SSc patients. Thus, the phase III trial of tocilizumab, which has been widely used in RA patients, showed effectiveness in stabilization and preservation of FVC, independent of the extent of radiographic ILD in lung from SSc patients, although tocilizumab failed to improve mRSS (88–90). Further investigation of these endotypes will determine how these groups respond to targeted interventions and whether they can predict which end organ would respond to a particular therapeutic.

---

**Supplemental Figure 1:**
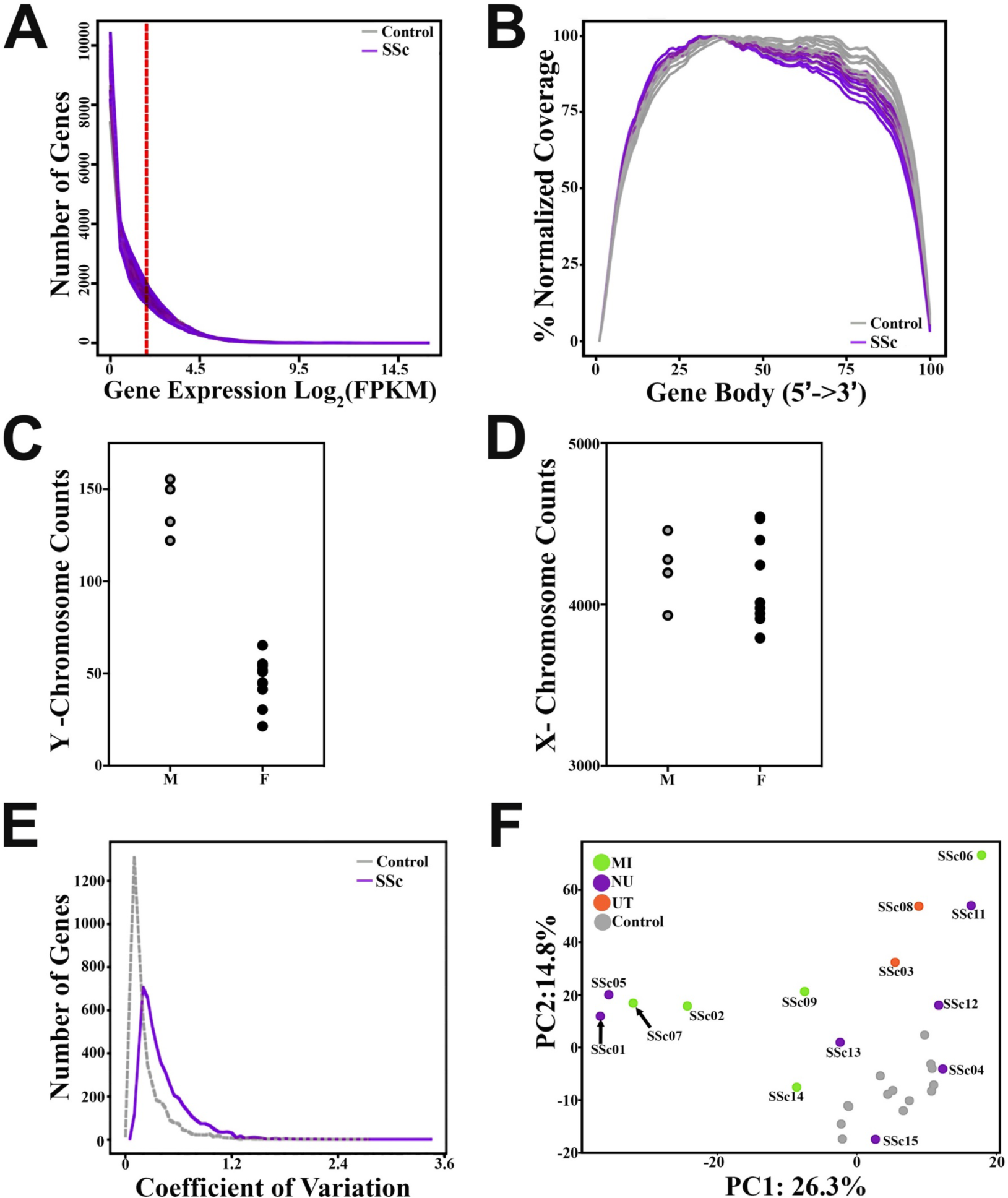
Quality Control analysis of Classical Monocyte RNA-seq. **(A)** Histogram indicating the number of genes expressed at the given Log2(FPKM) in each CM sample. Red line indicates the threshold for 5171 expressed genes (FPKM = 5) in at least 3 samples. **(B)** Normalized 5’-3’ gene coverage across length of genes. **(C)** Coefficient of variation histogram between control and SSc samples. **(D)** PCA of control and SSc samples are color coded based on sample collection sites. MI= University of Michigan. (green), NU, Northwestern University (purple), UT= University of Texas at Houston (orange). Sum of FPKM mapped to **(F)** Y- chromosome and **(G)** X-chromosome genes for each SSc CM sample.

**Supplemental Figure 2:**
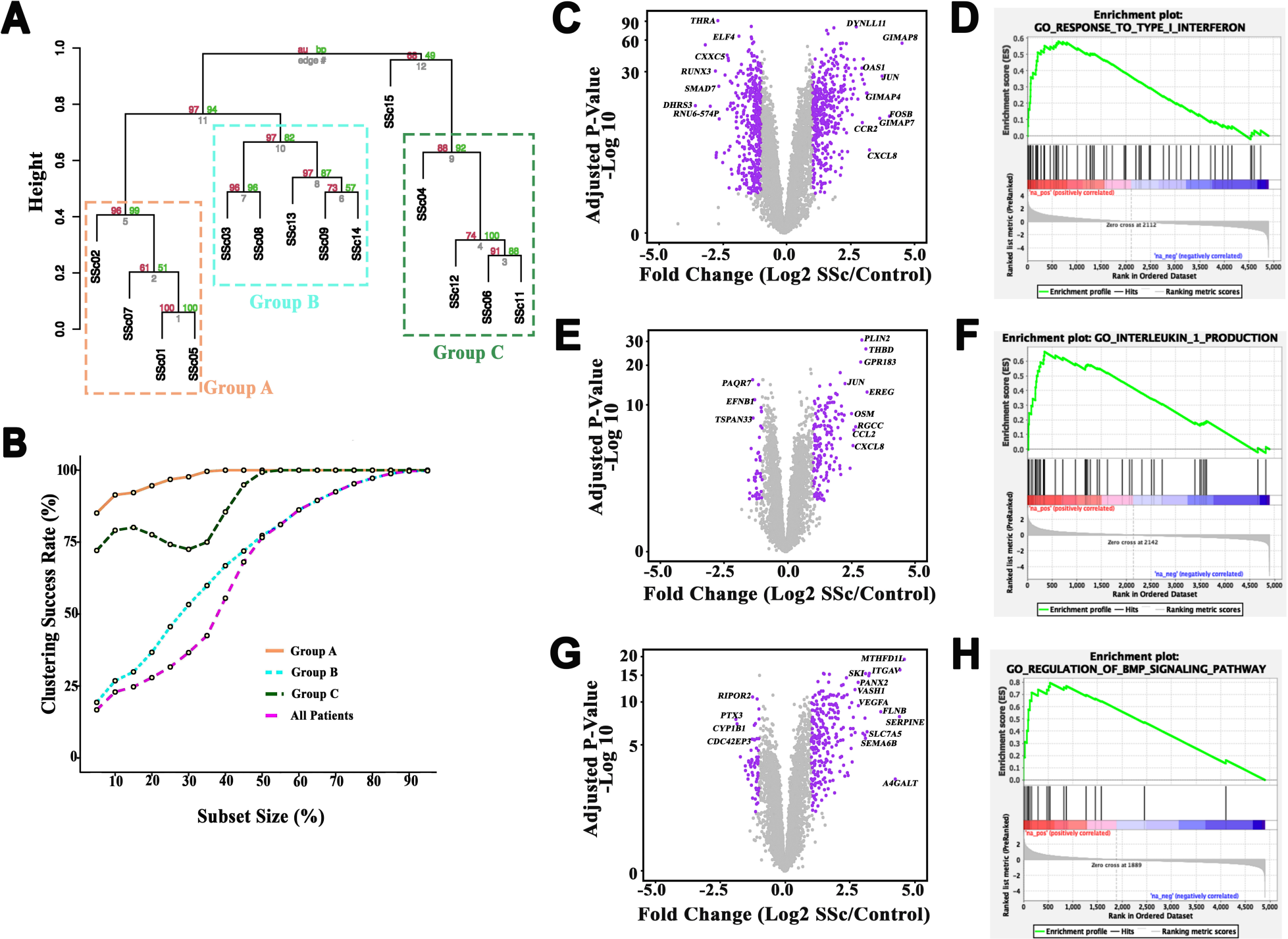
Differentially expressed genes vary by group in CM. Validation of the hierarchical clustering of CM gene expression in SSc patients. **(A)** Resampling of 1790 genes from 2A to calculate approximately unbiased (AU) probability (red) and bootstrap probability (BP) (green) of each edge (order in grey). **(B)** The percent of subset samples at various sizes that successfully recapitulate each patient group (A, B, C) or all patient groups. Volcano plot showing differential expression of genes in **(C)** Group A, 421 genes up regulated and 433 genes down regulated**(E)** Group B, 161 genes upregulated, and 47 genes down regulated and **(G)** Group C, 263 genes up regulated and 66 genes down regulated vs Control. Purple dots are Padj < 0.05 & log2 fold change >= 1 or <=-1. Representative enrichment plot from GSEA software for **(D)** Group A, **(F)** Group B and **(H)** Group C.

**Supplemental Figure 3:**
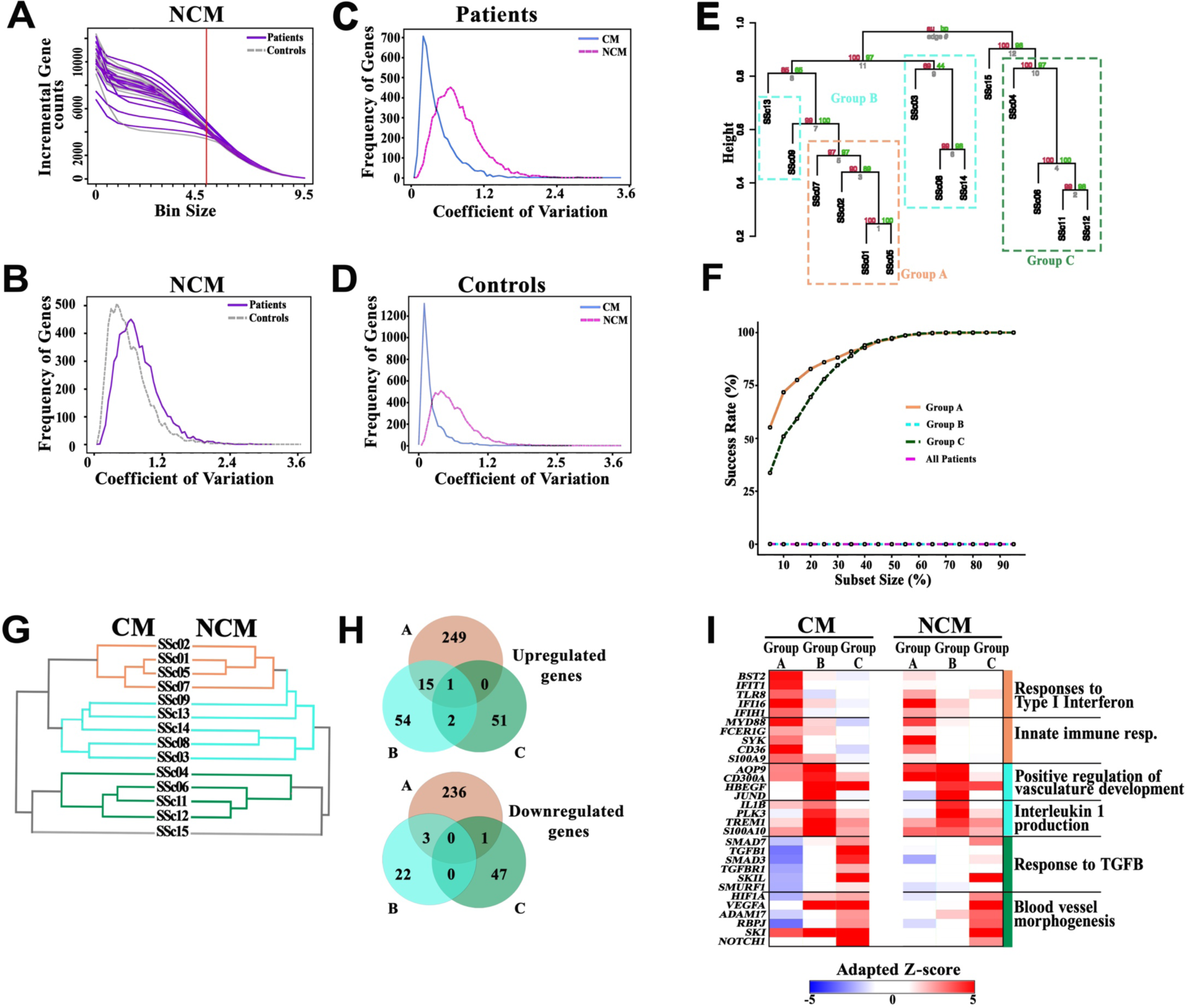
CM and NCM share transcriptional features in SSc patients. **(A)** Histogram indicating the number of genes expressed at the given Log2(CPM) in each NCM sample. Red line indicates the threshold for 7143 expressed genes (CPM = 31) in at least 3 samples. **(B)** Coefficient of variation histogram between control and SSc NCM samples. **(C-D)** Coefficient of variation histogram between CM and NCM of **(C)** SSc patients and **(D)** Controls. Validation of the hierarchical clustering of NCM gene expression in SSc patients. **(E)** Resampling of 1599 genes from 3A to calculate approximately unbiased (AU) probability (red) and bootstrap probability (BP) (green) of each edge (order in grey). **(F)** The percent of subset samples at various sizes that successfully recapitulate each patient group (A, B, C) or all patient groups. **(G)** Comparison of the hierarchical clustering dendrograms for SSc CM and NCM gene expression. **(H)** Venn Diagram showing overlap of upregulated (log2FC >1 and p-adj < 0.05) or downregulated (log2FC < -1 and p-adj < 0.05) genes in each patient group compared to controls. **(I)** Adapted z-score of relative expression in CM and NCM for representative genes associated with conserved processes patient groups.

**Supplemental Figure 4:**
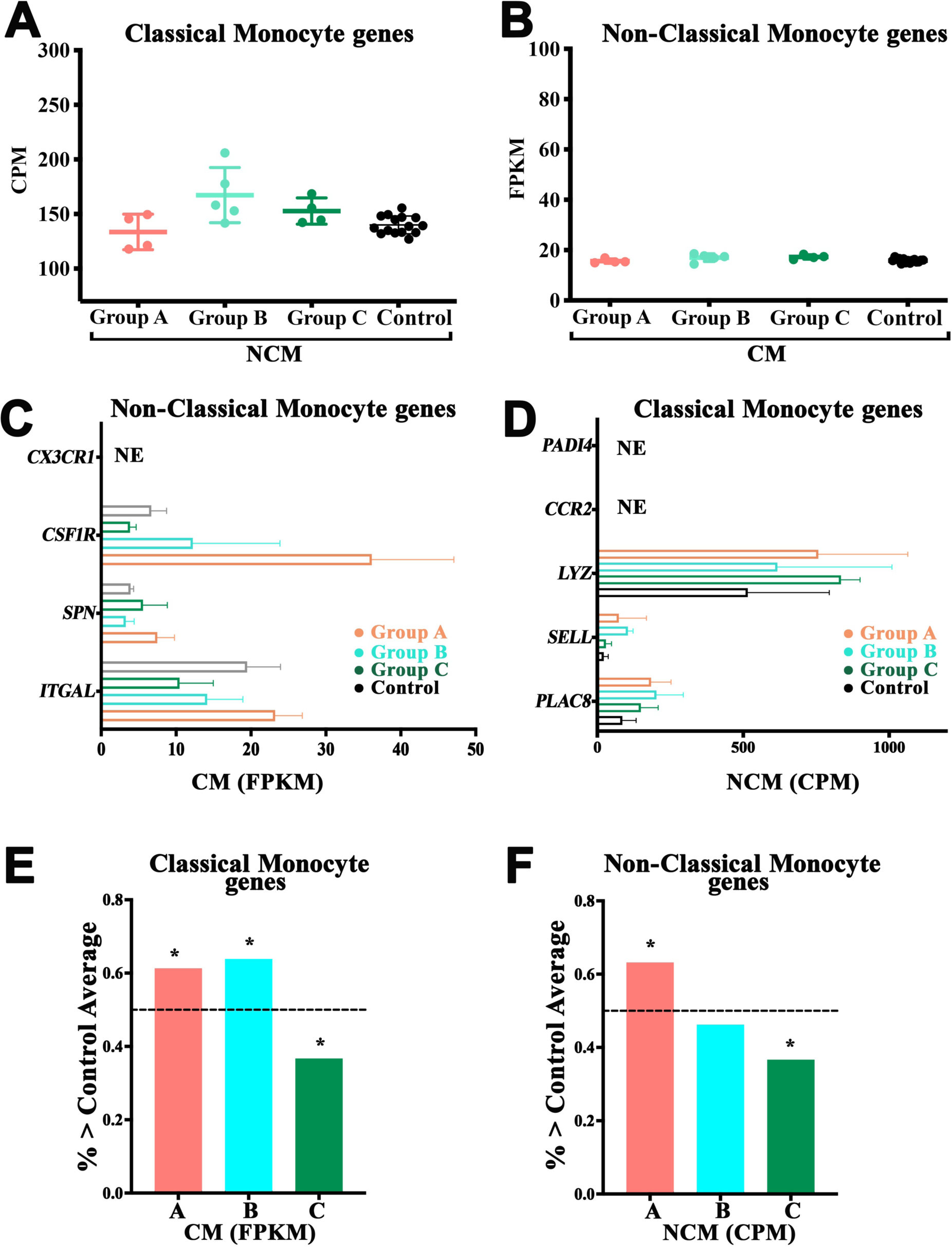
CM and NCM retain their individual cell type signature. **(A)** Average expression level of all CM genes in NCM of patients and controls **(B)** Average expression level of all NCM genes in CM of patients and controls. **(C)** Expression of representative NCM genes in patient CM samples by group. **(D)** Expression of representative CM genes in patient NCM samples by group. **E)** Percentage of CM genes in each patient CM sample by group that are expressed higher than control average. (**F)** Percentage of NCM genes in each patient NCM sample by group that are expressed higher than control average. * Indicates p < 0.5, Fisher’s exact test

**Supplemental Figure 5:**
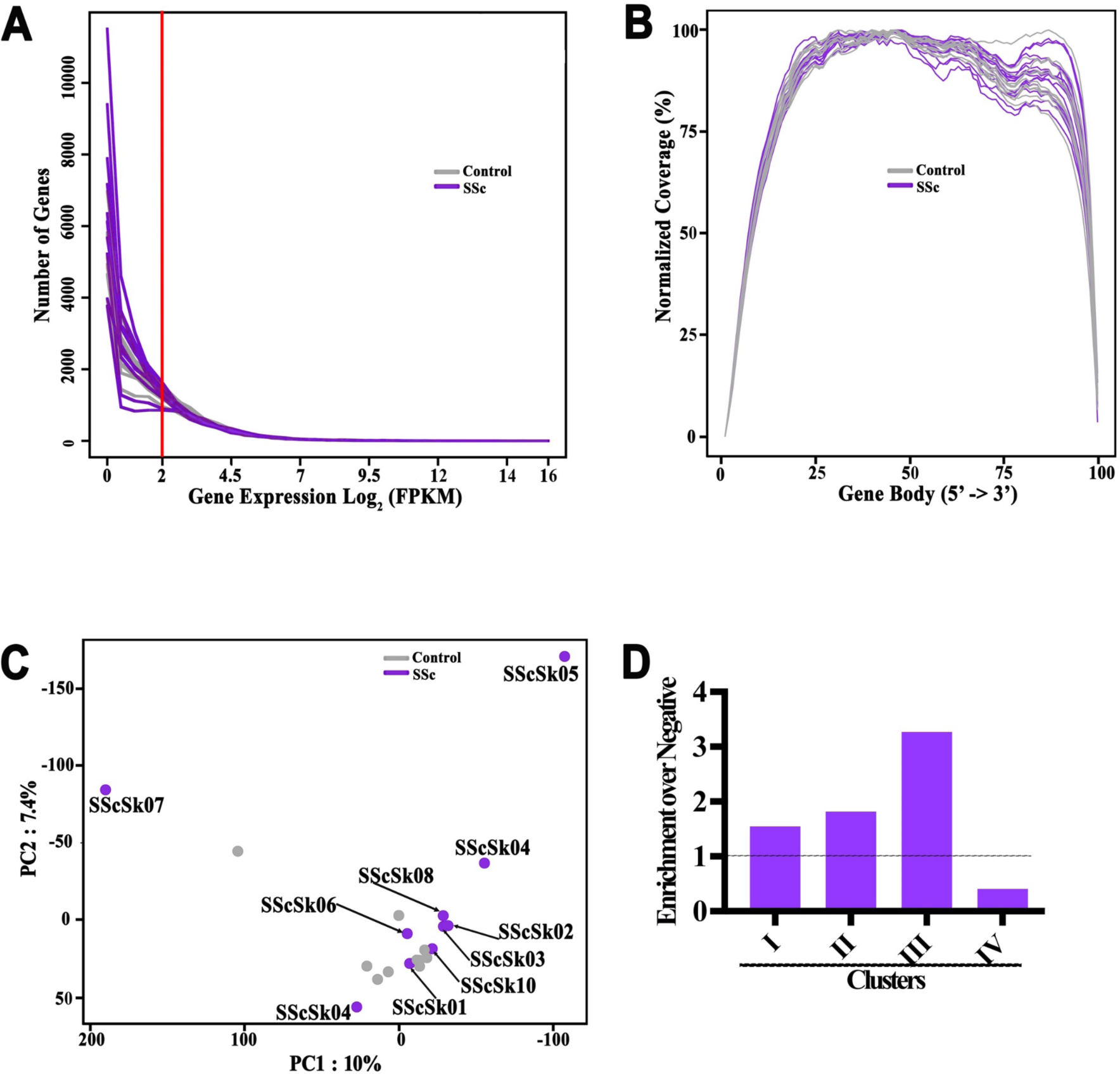
Gene expression in skin macrophages. **(A)** Histogram indicating the number of genes expressed at the given Log2(FPKM) in each skin macrophage sample. Red line indicates the threshold for 6339 expressed genes (FPKM = >3) in at least 3 samples. **(B)** 5’à3’ gene coverage in each patient and control sample. **(C)** PCA of skin macrophages samples annotated by SSc or control. **(D)** Bar graph showing ratio of genes from 5D that are enriched over negative relative to expected ratio (dotted line)

**Supplemental Figure 6:**
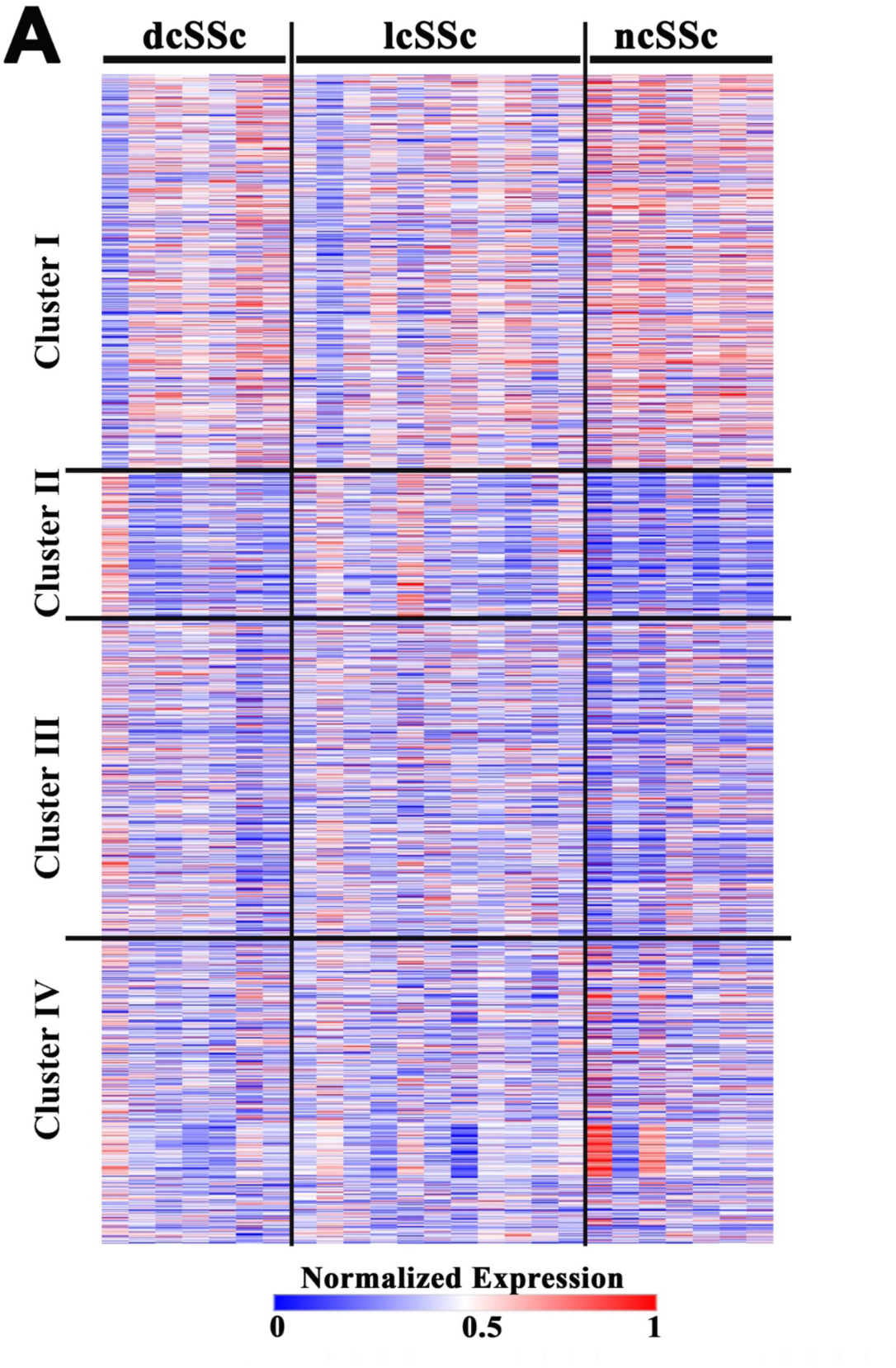
(A) Heatmap of normalized expression of genes per patient from CM Clusters I-IV (2A) in bulk monocytes from SSc patients categorized by disease phenotype (dataset described in van der Kroef, et al).

## References

1. Le EN, Wigley FM, Shah AA, Boin F, Hummers LK. Long-term experience of mycophenolate mofetil for treatment of diffuse cutaneous systemic sclerosis. Ann Rheum Dis. 2011.

2. Derk CT, Grace E, Shenin M, Naik M, Schulz S, Xiong W. A prospective open-label study of mycophenolate mofetil for the treatment of diffuse systemic sclerosis. Rheumatology (Oxford). 2009;48(12):1595–9.

3. Gerbino AJ, Goss CH, Molitor JA. Effect of mycophenolate mofetil on pulmonary function in scleroderma-associated interstitial lung disease. Chest. 2008;133(2):455–60.

4. Herrick AL, Lunt M, Whidby N, Ennis H, Silman A, McHugh N, et al. Observational study of treatment outcome in early diffuse cutaneous systemic sclerosis. J Rheumatol. 2010;37(1):116–24.

5. Liossis SN, Bounas A, Andonopoulos AP. Mycophenolate mofetil as first-line treatment improves clinically evident early scleroderma lung disease. Rheumatology (Oxford). 2006;45(8):1005–8.

6. Nihtyanova SI, Brough GM, Black CM, Denton CP. Mycophenolate mofetil in diffuse cutaneous systemic sclerosis--a retrospective analysis. Rheumatology (Oxford). 2007;46(3):442–5.

7. Plastiras SC, Vlachoyiannopoulos PG, Tzelepis GE. Mycophenolate mofetil for interstitial lung disease in scleroderma. Rheumatology (Oxford). 2006;45(12):1572.

8. Swigris JJ, Olson AL, Fischer A, Lynch DA, Cosgrove GP, Frankel SK, et al. Mycophenolate mofetil is safe, well tolerated, and preserves lung function in patients with connective tissue disease-related interstitial lung disease. Chest. 2006;130(1):30–6.

9. Vanthuyne M, Blockmans D, Westhovens R, Roufosse F, Cogan E, Coche E, et al. A pilot study of mycophenolate mofetil combined to intravenous methylprednisolone pulses and oral low-dose glucocorticoids in severe early systemic sclerosis. Clin Exp Rheumatol. 2007;25(2):287–92.

10. Zamora AC, Wolters PJ, Collard HR, Connolly MK, Elicker BM, Webb WR, et al. Use of mycophenolate mofetil to treat scleroderma-associated interstitial lung disease. Respir Med. 2008;102(1):150–5.

11. Fischer A, Brown KK, Du Bois RM, Frankel SK, Cosgrove GP, Fernandez-Perez ER, et al. Mycophenolate mofetil improves lung function in connective tissue disease-associated interstitial lung disease. J Rheumatol. 2013;40(5):640–6.

12. Assassi S, Radstake TR, Mayes MD, Martin J. Genetics of scleroderma: implications for personalized medicine? BMC Med. 2013;11:9.

13. Prescott RJ, Freemont AJ, Jones CJ, Hoyland J, Fielding P. Sequential dermal microvascular and perivascular changes in the development of scleroderma. The Journal of pathology. 1992;166(3):255–63.

14. Cossu M, van Bon L, Preti C, Rossato M, Beretta L, Radstake T. Earliest Phase of Systemic Sclerosis Typified by Increased Levels of Inflammatory Proteins in the Serum. Arthritis Rheumatol. 2017;69(12):2359–69.

15. Lescoat A, Lecureur V, Roussel M, Sunnaram BL, Ballerie A, Coiffier G, et al. CD16- positive circulating monocytes and fibrotic manifestations of systemic sclerosis. Clin Rheumatol. 2017;36(7):1649–54.

16. Vreca M, Zekovic A, Damjanov N, Andjelkovic M, Ugrin M, Pavlovic S, et al. Expression of TLR7, TLR9, JAK2, and STAT3 genes in peripheral blood mononuclear cells from patients with systemic sclerosis. J Appl Genet. 2018;59(1):59–66.

17. Burt RK, Shah SJ, Dill K, Grant T, Gheorghiade M, Schroeder J, et al. Autologous non-myeloablative haemopoietic stem-cell transplantation compared with pulse cyclophosphamide once per month for systemic sclerosis (ASSIST): an open-label, randomised phase 2 trial. Lancet. 2011;378(9790):498–506.

18. Rezaei R, Mahmoudi M, Gharibdoost F, Kavosi H, Dashti N, Imeni V, et al. IRF7 gene expression profile and methylation of its promoter region in patients with systemic sclerosis. Int J Rheum Dis. 2017;20(10):1551–61.

19. van Laar JM, Farge D, Sont JK, Naraghi K, Marjanovic Z, Larghero J, et al. Autologous hematopoietic stem cell transplantation vs intravenous pulse cyclophosphamide in diffuse cutaneous systemic sclerosis: a randomized clinical trial. JAMA. 2014;311(24):2490–8.

20. Ziegler-Heitbrock L, Ancuta P, Crowe S, Dalod M, Grau V, Hart DN, et al. Nomenclature of monocytes and dendritic cells in blood. Blood. 2010;116(16):e74–80.

21. Yona S, Jung S. Monocytes: subsets, origins, fates and functions. Current Opinion in Hematology. 2010;17(1):53–9.

22. van der Kroef M, van den Hoogen LL, Mertens JS, Blokland SLM, Haskett S, Devaprasad A, et al. Cytometry by time of flight identifies distinct signatures in patients with systemic sclerosis, systemic lupus erythematosus and Sjogrens syndrome. Eur J Immunol. 2020;50(1):119–29.

23. Toledo DM, Pioli PA. Macrophages in Systemic Sclerosis: Novel Insights and Therapeutic Implications. Current rheumatology reports. 2019;21(7):31.

24. Higashi-Kuwata N, Jinnin M, Makino T, Fukushima S, Inoue Y, Muchemwa FC, et al. Characterization of monocyte/macrophage subsets in the skin and peripheral blood derived from patients with systemic sclerosis. Arthritis Res Ther. 2010;12(4):R128.

25. Misharin AV, Morales-Nebreda L, Reyfman PA, Cuda CM, Walter JM, McQuattie-Pimentel AC, et al. Monocyte-derived alveolar macrophages drive lung fibrosis and persist in the lung over the life span. J Exp Med. 2017;214(8):2387–404.

26. Reyfman PA, Walter JM, Joshi N, Anekalla KR, McQuattie-Pimentel AC, Chiu S, et al. Single-Cell Transcriptomic Analysis of Human Lung Provides Insights into the Pathobiology of Pulmonary Fibrosis. Am J Respir Crit Care Med. 2019;199(12):1517–36.

27. Mathai SK, Gulati M, Peng X, Russell TR, Shaw AC, Rubinowitz AN, et al. Circulating monocytes from systemic sclerosis patients with interstitial lung disease show an enhanced profibrotic phenotype. Laboratory investigation; a journal of technical methods and pathology. 2010;90(6):812–23.

28. Arai M, Ikawa Y, Chujo S, Hamaguchi Y, Ishida W, Shirasaki F, et al. Chemokine receptors CCR2 and CX3CR1 regulate skin fibrosis in the mouse model of cytokine-induced systemic sclerosis. J Dermatol Sci. 2013;69(3):250–8.

29. Hinchcliff M, Toledo DM, Taroni JN, Wood TA, Franks JM, Ball MS, et al. Mycophenolate Mofetil Treatment of Systemic Sclerosis Reduces Myeloid Cell Numbers and Attenuates the Inflammatory Gene Signature in Skin. J Invest Dermatol. 2018.

30. Bossini-Castillo L, Villanueva-Martin G, Kerick M, Acosta-Herrera M, Lopez-Isac E, Simeon CP, et al. Genomic Risk Score impact on susceptibility to systemic sclerosis. Ann Rheum Dis. 2021;80(1):118–27.

31. Beretta L, Barturen G, Vigone B, Bellocchi C, Hunzelmann N, De Langhe E, et al. Genome-wide whole blood transcriptome profiling in a large European cohort of systemic sclerosis patients. Ann Rheum Dis. 2020;79(9):1218–26.

32. Milano A, Pendergrass SA, Sargent JL, George LK, McCalmont TH, Connolly MK, et al. Molecular subsets in the gene expression signatures of scleroderma skin. PLoS One. 2008;3(7):e2696.

33. Pendergrass SA, Lemaire R, Francis IP, Mahoney JM, Lafyatis R, Whitfield ML. Intrinsic gene expression subsets of diffuse cutaneous systemic sclerosis are stable in serial skin biopsies. J Invest Dermatol. 2012;132(5):1363–73.

34. Taroni JN, Martyanov V, Huang CC, Mahoney JM, Hirano I, Shetuni B, et al. Molecular characterization of systemic sclerosis esophageal pathology identifies inflammatory and proliferative signatures. Arthritis Res Ther. 2015;17:194.

35. Franks JM, Martyanov V, Wang Y, Wood TA, Pinckney A, Crofford LJ, et al. Machine learning predicts stem cell transplant response in severe scleroderma. Ann Rheum Dis. 2020;79(12):1608–15.

36. Assassi S, Swindell WR, Wu M, Tan FD, Khanna D, Furst DE, et al. Dissecting the heterogeneity of skin gene expression patterns in systemic sclerosis. Arthritis Rheumatol. 2015;67(11):3016–26.

37. Hinchcliff M, Huang CC, Wood TA, Matthew Mahoney J, Martyanov V, Bhattacharyya S, et al. Molecular signatures in skin associated with clinical improvement during mycophenolate treatment in systemic sclerosis. J Invest Dermatol. 2013;133(8):1979–89.

38. Chakravarty EF, Martyanov V, Fiorentino D, Wood TA, Haddon DJ, Jarrell JA, et al. Gene expression changes reflect clinical response in a placebo-controlled randomized trial of abatacept in patients with diffuse cutaneous systemic sclerosis. Arthritis Res Ther. 2015;17:159.

39. Assassi S, Li N, Volkmann ER, Mayes MD, Runger D, Ying J, et al. Predictive Significance of Serum Interferon-Inducible Protein Score for Response to Treatment in Systemic Sclerosis-Related Interstitial Lung Disease. Arthritis Rheumatol. 2021;73(6):1005–13.

40. Bernstein EJ, Jaafar S, Assassi S, Domsic RT, Frech TM, Gordon JK, et al. Performance Characteristics of Pulmonary Function Tests for the Detection of Interstitial Lung Disease in Adults With Early Diffuse Cutaneous Systemic Sclerosis. Arthritis Rheumatol. 2020;72(11):1892–6.

41. Frech TM, Revelo MP, Ryan JJ, Shah AA, Gordon J, Domsic R, et al. Cardiac metabolomics and autopsy in a patient with early diffuse systemic sclerosis presenting with dyspnea: a case report. J Med Case Rep. 2015;9:136.

42. Gordon JK, Girish G, Berrocal VJ, Zhang M, Hatzis C, Assassi S, et al. Reliability and Validity of the Tender and Swollen Joint Counts and the Modified Rodnan Skin Score in Early Diffuse Cutaneous Systemic Sclerosis: Analysis from the Prospective Registry of Early Systemic Sclerosis Cohort. J Rheumatol. 2017;44(6):791–4.

43. Jaafar S, Lescoat A, Huang S, Gordon J, Hinchcliff M, Shah AA, et al. Clinical characteristics, visceral involvement, and mortality in at-risk or early diffuse systemic sclerosis: a longitudinal analysis of an observational prospective multicenter US cohort. Arthritis Res Ther. 2021;23(1):170.

44. Skaug B, Khanna D, Swindell WR, Hinchcliff ME, Frech TM, Steen VD, et al. Global skin gene expression analysis of early diffuse cutaneous systemic sclerosis shows a prominent innate and adaptive inflammatory profile. Ann Rheum Dis. 2020;79(3):379–86.

45. Mandelin AM, 2nd, Homan PJ, Shaffer AM, Cuda CM, Dominguez ST, Bacalao E, et al. Transcriptional Profiling of Synovial Macrophages Using Minimally Invasive Ultrasound-Guided Synovial Biopsies in Rheumatoid Arthritis. Arthritis Rheumatol. 2018;70(6):841–54.

46. Kim D, Pertea G, Trapnell C, Pimentel H, Kelley R, Salzberg SL. TopHat2: accurate alignment of transcriptomes in the presence of insertions, deletions and gene fusions. Genome Biol. 2013;14(4):R36.

47. Dobin A, Davis CA, Schlesinger F, Drenkow J, Zaleski C, Jha S, et al. STAR: ultrafast universal RNA-seq aligner. Bioinformatics. 2013;29(1):15–21.

48. Anders S, Pyl PT, Huber W. HTSeq--a Python framework to work with high-throughput sequencing data. Bioinformatics. 2015;31(2):166–9.

49. Wang L, Wang S, Li W. RSeQC: quality control of RNA-seq experiments. Bioinformatics. 2012;28(16):2184–5.

50. Love MI, Huber W, Anders S. Moderated estimation of fold change and dispersion for RNA-seq data with DESeq2. Genome Biol. 2014;15(12):550.

51. Suzuki R, Shimodaira H. Pvclust: an R package for assessing the uncertainty in hierarchical clustering. Bioinformatics. 2006;22(12):1540–2.

52. Eden E, Navon R, Steinfeld I, Lipson D, Yakhini Z. GOrilla: a tool for discovery and visualization of enriched GO terms in ranked gene lists. BMC Bioinformatics. 2009;10:48.

53. Wong KL, Tai JJ, Wong WC, Han H, Sem X, Yeap WH, et al. Gene expression profiling reveals the defining features of the classical, intermediate, and nonclassical human monocyte subsets. Blood. 2011;118(5):e16–31.

54. Robin X, Turck N, Hainard A, Tiberti N, Lisacek F, Sanchez JC, et al. pROC: an open-source package for R and S+ to analyze and compare ROC curves. BMC Bioinformatics. 2011;12:77.

55. Mason S.J., GrahamN. E. Areas beneath the relative operating characteristics (ROC) and relative operating levels (ROL) curves: Statistical significance and interpretation, . Q J R Meteorol Soc 2002;128: 2145--66.

56. van der Kroef M, Castellucci M, Mokry M, Cossu M, Garonzi M, Bossini-Castillo LM, et al. Histone modifications underlie monocyte dysregulation in patients with systemic sclerosis, underlining the treatment potential of epigenetic targeting. Ann Rheum Dis. 2019;78(4):529–38.

57. Mehta BK, Espinoza ME, Hinchcliff M, Whitfield ML. Molecular “omic” signatures in systemic sclerosis. Eur J Rheumatol. 2020;7(Suppl 3):S173–S80.

58. Dao DT, Anez-Bustillos L, Adam RM, Puder M, Bielenberg DR. Heparin-Binding Epidermal Growth Factor-Like Growth Factor as a Critical Mediator of Tissue Repair and Regeneration. Am J Pathol. 2018;188(11):2446–56.

59. Salcedo R, Oppenheim JJ. Role of chemokines in angiogenesis: CXCL12/SDF-1 and CXCR4 interaction, a key regulator of endothelial cell responses. Microcirculation. 2003;10(3- 4):359–70.

60. Chandel S, Manikandan A, Mehta N, Nathan AA, Tiwari RK, Mohapatra SB, et al. The protein tyrosine phosphatase PTP-PEST mediates hypoxia-induced endothelial autophagy and angiogenesis via AMPK activation. J Cell Sci. 2021;134(1).

61. Yona S, Kim KW, Wolf Y, Mildner A, Varol D, Breker M, et al. Fate mapping reveals origins and dynamics of monocytes and tissue macrophages under homeostasis. Immunity. 2013;38(1):79–91.

62. Guilliams M, Mildner A, Yona S. Developmental and Functional Heterogeneity of Monocytes. 2018;49(4):595–613.

63. Mildner A, Schonheit J, Giladi A, David E, Lara-Astiaso D, Lorenzo-Vivas E, et al. Genomic Characterization of Murine Monocytes Reveals C/EBPbeta Transcription Factor Dependence of Ly6C(-) Cells. Immunity. 2017;46(5):849–62 e7.

64. Kapellos TS, Bonaguro L, Gemund I, Reusch N, Saglam A, Hinkley ER, et al. Human Monocyte Subsets and Phenotypes in Major Chronic Inflammatory Diseases. Front Immunol. 2019;10:2035.

65. Watanabe S, Alexander M, Misharin AV, Budinger GRS. The role of macrophages in the resolution of inflammation. J Clin Invest. 2019;129(7):2619–28.

66. Ainola M, Li TF, Mandelin J, Hukkanen M, Choi SJ, Salo J, et al. Involvement of a disintegrin and a metalloproteinase 8 (ADAM8) in osteoclastogenesis and pathological bone destruction. Ann Rheum Dis. 2009;68(3):427–34.

67. Snodgrass RG, Brune B. Regulation and Functions of 15-Lipoxygenases in Human Macrophages. Front Pharmacol. 2019;10:719.

68. Hoober JK, Eggink LL, Cote R. Stories From the Dendritic Cell Guardhouse. Front Immunol. 2019;10:2880.

69. Dorrington MG, Fraser IDC. NF-kappaB Signaling in Macrophages: Dynamics, Crosstalk, and Signal Integration. Front Immunol. 2019;10:705.

70. Shen YM, Zhao Y, Zeng Y, Yan L, Chen BL, Leng AM, et al. Inhibition of Pim-1 kinase ameliorates dextran sodium sulfate-induced colitis in mice. Dig Dis Sci. 2012;57(7):1822–31.

71. Ko CY, Chang WC, Wang JM. Biological roles of CCAAT/Enhancer-binding protein delta during inflammation. J Biomed Sci. 2015;22:6.

72. Chang LH, Huang HS, Wu PT, Jou IM, Pan MH, Chang WC, et al. Role of macrophage CCAAT/enhancer binding protein delta in the pathogenesis of rheumatoid arthritis in collagen-induced arthritic mice. PLoS One. 2012;7(9):e45378.

73. Mass E, Ballesteros I, Farlik M, Halbritter F, Gunther P, Crozet L, et al. Specification of tissue-resident macrophages during organogenesis. Science. 2016;353(6304).

74. Li X, Zhang X, Xia J, Zhang L, Chen B, Lian G, et al. Macrophage HIF-2alpha suppresses NLRP3 inflammasome activation and alleviates insulin resistance. Cell Rep. 2021;36(8):109607.

75. Wang X, Abraham S, McKenzie JAG, Jeffs N, Swire M, Tripathi VB, et al. LRG1 promotes angiogenesis by modulating endothelial TGF-beta signalling. Nature. 2013;499(7458):306–11.

76. Villani AC, Satija R, Reynolds G, Sarkizova S, Shekhar K, Fletcher J, et al. Single-cell RNA-seq reveals new types of human blood dendritic cells, monocytes, and progenitors. Science. 2017;356(6335).

77. Dutertre CA, Becht E, Irac SE, Khalilnezhad A, Narang V, Khalilnezhad S, et al. Single-Cell Analysis of Human Mononuclear Phagocytes Reveals Subset-Defining Markers and Identifies Circulating Inflammatory Dendritic Cells. Immunity. 2019;51(3):573–89 e8.

78. Higgs BW, Liu Z, White B, Zhu W, White WI, Morehouse C, et al. Patients with systemic lupus erythematosus, myositis, rheumatoid arthritis and scleroderma share activation of a common type I interferon pathway. Ann Rheum Dis. 2011;70(11):2029–36.

79. Johnson ME, Mahoney JM, Taroni J, Sargent JL, Marmarelis E, Wu MR, et al. Experimentally-derived fibroblast gene signatures identify molecular pathways associated with distinct subsets of systemic sclerosis patients in three independent cohorts. PLoS One. 2015;10(1):e0114017.

80. Taroni JN, Greene CS, Martyanov V, Wood TA, Christmann RB, Farber HW, et al. A novel multi-network approach reveals tissue-specific cellular modulators of fibrosis in systemic sclerosis. Genome Med. 2017;9(1):27.

81. Mahoney JM, Taroni J, Martyanov V, Wood TA, Greene CS, Pioli PA, et al. Systems level analysis of systemic sclerosis shows a network of immune and profibrotic pathways connected with genetic polymorphisms. PLoS Comput Biol. 2015;11(1):e1004005.

82. Showalter K, Spiera R, Magro C, Agius P, Martyanov V, Franks JM, et al. Machine learning integration of scleroderma histology and gene expression identifies fibroblast polarisation as a hallmark of clinical severity and improvement. Ann Rheum Dis. 2021;80(2):228–37.

83. Skaug B, Lyons MA, Swindell WR, Salazar GA, Wu M, Tran TM, et al. Large-scale analysis of longitudinal skin gene expression in systemic sclerosis reveals relationships of immune cell and fibroblast activity with skin thickness and a trend towards normalisation over time. Ann Rheum Dis. 2021.

84. Lofgren S, Hinchcliff M, Carns M, Wood T, Aren K, Arroyo E, et al. Integrated, multicohort analysis of systemic sclerosis identifies robust transcriptional signature of disease severity. JCI Insight. 2016;1(21):e89073.

85. Kobayashi S, Nagafuchi Y, Okubo M, Sugimori Y, Shirai H, Hatano H, et al. Integrated bulk and single-cell RNA-sequencing identified disease-relevant monocytes and a gene network module underlying systemic sclerosis. J Autoimmun. 2021;116:102547.

86. Xue D, Tabib T, Morse C, Yang Y, Domsic R, Khanna D, et al. Expansion of FCGR3A(+) macrophages, FCN1(+) mo-DC, and plasmacytoid dendritic cells associated with severe skin disease in systemic sclerosis. Arthritis Rheumatol. 2021.

87. Jaeger VK, Wirz EG, Allanore Y, Rossbach P, Riemekasten G, Hachulla E, et al. Incidences and Risk Factors of Organ Manifestations in the Early Course of Systemic Sclerosis: A Longitudinal EUSTAR Study. PLoS One. 2016;11(10):e0163894.

88. Suleman Y, Clark KEN, Cole AR, Ong VH, Denton CP. Real-world experience of Tocilizumab in systemic sclerosis: potential benefit on lung function for anti-topoisomerase (ATA) positive patients. Rheumatology (Oxford). 2021.

89. Roofeh D, Lin CJF, Goldin J, Kim GH, Furst DE, Denton CP, et al. Tocilizumab Prevents Progression of Early Systemic Sclerosis Associated Interstitial Lung Disease. Arthritis Rheumatol. 2021.

90. Khanna D, Lin CJF, Furst DE, Goldin J, Kim G, Kuwana M, et al. Tocilizumab in systemic sclerosis: a randomised, double-blind, placebo-controlled, phase 3 trial. Lancet Respir Med. 2020;8(10):963–74.

91. Khanna D, Denton CP, Lin CJF, van Laar JM, Frech TM, Anderson ME, et al. Safety and efficacy of subcutaneous tocilizumab in systemic sclerosis: results from the open-label period of a phase II randomised controlled trial (faSScinate). Ann Rheum Dis. 2018;77(2):212–20.

92. Khanna D, Denton CP, Jahreis A, van Laar JM, Frech TM, Anderson ME, et al. Safety and efficacy of subcutaneous tocilizumab in adults with systemic sclerosis (faSScinate): a phase 2, randomised, controlled trial. Lancet. 2016;387(10038):2630–40.

93. Arnold MB, Khanna D, Denton CP, van Laar JM, Frech TM, Anderson ME, et al. Patient acceptable symptom state in scleroderma: results from the tocilizumab compared with placebo trial in active diffuse cutaneous systemic sclerosis. Rheumatology (Oxford). 2018;57(1):152–7.

